# Comprehensive characterization of migration profiles of murine cerebral cortical neurons during development using FlashTag labeling

**DOI:** 10.1101/2020.10.05.317925

**Authors:** Satoshi Yoshinaga, Minkyung Shin, Ayako Kitazawa, Kazuhiro Ishii, Masato Tanuma, Atsushi Kasai, Hitoshi Hashimoto, Ken-ichiro Kubo, Kazunori Nakajima

## Abstract

In mammalian cerebral neocortex, different regions have different cytoarchitecture, neuronal birthdates and functions. In most regions, neuronal migratory profiles have been speculated similar to each other based on observations using thymidine analogues. Few reports investigated regional migratory differences from mitosis at the ventricular surface. Here, in mice, we applied FlashTag technology, in which dyes are injected intraventricularly, to describe migratory profiles. We revealed a mediolateral regional difference in migratory profiles of neurons that is dependent on the developmental stages, e.g., neurons labeled at E12.5-15.5 reached their destination earlier dorsomedially than dorsolaterally even where there were underlying ventricular surfaces, reflecting sojourning below the subplate. This difference was hardly recapitulated by thymidine analogues, which visualize neurogenic gradient, suggesting biological significance different from neurogenic gradient. These observations advance understanding of cortical development, portraying strength of FlashTag in studying migration, and are thus a resource for studies of normal and abnormal neurodevelopment.

## Introduction

The mammalian cerebral neocortex is a well-organized, 6-layered structure that contains diversity of neurons. Neuronal migration is an essential step for precise formation of complex cortical cytoarchitecture which underlies the evolution of mammalian cognitive function. In the earliest stage of cortical development, neural stem cells form a pseudostratified structure called neuroepithelium, and these stem cells undergo self-renewal to expand the cortical areas (Caviness et al., 1995; His, 1889; Rakic, 1995; Sauer, 1935; Subramanian et al., 2017). They then begin to produce the earliest-born neurons (Bystron et al., 2006; Iacopetti et al., 1999), which form the preplate (PP), or primordial plexiform layer (Marin-Padilla, 1971) (See Figures S1A, B and the *histological terminology* section in Star Methods). These neurons include Cajal-Retzius cells and future subplate (SP) neurons, most of which are transient population that undergo cell death postnatally (Hoerder-Suabedissen and Molnár, 2015; Kostovic and Rakic, 1990; Price et al., 1997). In the pallium, cortical projection neuron production follows. They derive from radial glial cells in the ventricular zone (VZ). Some daughter cells become postmitotic soon after they exit the VZ (Tabata et al., 2009) and others divide in a more basal structure (the subventricular zone or SVZ)(Boulder-Committee, 1970; Takahashi et al., 1996). In both cases, they migrate radially through the intermediate zone (IZ), SP, and the cortical plate (CP) to the primitive cortical zone (PCZ) (Sekine et al., 2011; Sekine et al., 2012), the most superficial part of the CP. Migrating neurons overtake earlier born neurons to finish their migration in the PCZ and this process serves as a basis of the inside-out pattern of neuronal positioning, in which earlier-born neurons position deeply and later-born neurons position superficially (Sekine et al., 2011; Shin et al., 2019).

Different cortical regions have different functions. The cerebral cortex is subdivided into many cortical areas based on cytoarchitectonics (Brodmann, 1909), which have high correlations with function. According to the protomap hypothesis, the neural stem cells in the VZ provide a protomap of prospective cytoarchitectonic areas (Rakic, 1988). Since, in addition to thalamic input (Moreno-Juan et al., 2017), neuronal migration takes place between proliferation at the ventricular surface and formation of cytoarchitectonics, neuronal migratory profiles of the whole brain visualized from mitosis at the ventricular surface to their final destination may serve as basic information to understand the formation of the complex mammalian brain.

To describe the above-mentioned neuronal behaviors in the neurogenesis, positioning and neuronal migration, thymidine analogues have long been used. Interkinetic nuclear migration of the VZ stem cells (Fujita, 1963) and inside-out pattern of neuronal birthdate (Angevine and Sidman, 1961; Bayer and Altman, 1991; Hicks and D’Amato, 1968) was clearly shown by tritium thymidine (^3^H-TdR). The limitation of the use of these S-phase markers to study neuronal migration was hardly discussed, however; in the last 20 years, growing evidence suggests that many projection neurons, especially superficial layer neurons, are generated indirectly in the SVZ from intermediate neural progenitors (Haubensak et al., 2004; Kowalczyk et al., 2009; Miyata et al., 2004; Noctor et al., 2004; Takahashi et al., 1996) and basal radial glial cells (Fietz et al., 2010; Hansen et al., 2010) that derive from (apical) radial glial cells, in addition to direct neurogenesis (Tabata et al., 2009). Also, interneurons are born in the ventral forebrain and migrate all the way to the cortex (tangential migration) (Anderson et al., 1997; Marin and Rubenstein, 2001; Tamamaki et al., 1997). Thymidine analogues are incorporated in the S-phase and retained by the progeny of dividing cells that undergo final mitosis irrespective of the anatomical position. Because basal progenitors are already in the midst of migration when they are in the S-phase, and because interneurons incorporate thymidine analogues ventrally, migratory “profiles” of neurons revealed by thymidine analogues contain those with different “starting points” (cellular positions when thymidine analogues are incorporated).

As a method to align the “starting points”, we previously developed *in utero* electroporation, in which expression plasmids are injected into the lateral ventricle and electrical pulses are given to transfect the cells along the ventricular surface at the time of labeling (Tabata and Nakajima, 2001; Tabata and Nakajima, 2008). This method is supposed to label apical progenitors in the S/G2/M phase preferentially (Pilaz et al., 2009). With this method, we previously described a) different migratory profiles between the direct progeny of apical progenitors and basal progenitors and b) regional differences in the abundance of the two modes of neurogenesis between dorsomedial and dorsolateral cortex (Tabata et al., 2009). Since it is well described that neurogenic event progresses along the lateral-to-medial gradient (Hicks and D’Amato, 1968; Smart and McSherry, 1982; Smart and Smart, 1982; Takahashi et al., 1999), we hypothesized that the migratory profile of dorsomedial (future cingulate) cortex and dorsolateral cortex (future somatosensory cortex where there is an underlying ventricular zone prenatally) differs significantly. On the other hand, aforementioned work (Bayer and Altman, 1991) and others (Hicks and D’Amato, 1968) using thymidine analogues did not describe as such, although the former studied regional differences in the migration of later-born cortical neurons in rats and they observed that it took about two days for labeled neurons to reach the top of the CP in the dorsomedial and dorsolateral cortex where there is an underlining VZ, while neurons migrating to the lateral (future presumptive insular and piriform) cortex which lack underlining VZ took longer because they migrate in a sigmoid manner to circumvent the growing striatum along the lateral cortical stream. The overall migratory profiles of neurons that are born at ventricular surfaces in different cortical regions at different stages remain to be described.

To visualize migration of neurons of different cortical regions that undergo mitosis at ventricular surface at given timing, we decided to take advantage of FlashTag (FT) technology (Govindan et al., 2018; Telley et al., 2016), in which fluorescent dyes were injected into the ventricle. This technique is reported to label ventricular cells at the M-phase specifically. Once the hydrophobic precursor fluorescent molecules (5(6)-carboxyfluorescein diacetate succinimidyl ester; CFDA-SE, often called CFSE in biological contexts) diffuse into the cell, cellular esterases cleave it to produce carboxyfluorescein succinimidyl ester, which is fluorescent and covalently bound to intracellular proteins (Telley et al., 2016). FT also refers to the use of other compounds with identical modes of action, including CytoTell Blue (Telley et al., 2016). Here, we successfully visualized the migration of projection neuron in different cortical regions with high temporal resolution. We describe mediolateral regional differences in migratory profiles of neurons born at different stages in regions where there is an underlying ventricular zone, which was not clearly detected by experiments using thymidine analogues.

## Results

### Characterization of cell population labeled with FT

FT technology has been increasingly used to label neuronal progenitors on the ventricular surfaces and their positioning (Govindan et al., 2018; Mayer et al., 2018; Oberst et al., 2019; Telley et al., 2019; Telley et al., 2016), but it has hardly used to study overall migratory profiles. Therefore, we first characterized the cellular population labeled with FT to ensure the validity of our analyses. We performed intraventricular injection of ∼0.5 μl of 1 mM of CFSE. We decreased the concentration of CFSE and quantity of organic solvent to one tenth of that used in the previous studies (Govindan et al., 2018; Telley et al., 2016) to minimize any possible but unknown side effects from injection although the original concentration was successfully used. To characterize the population of cells labeled with FT, we injected CFSE into the lateral ventricle (LV) of E14 ICR embryos and fixed 0.5-9.5 hours later (Figures 1A-E). When brains were fixed 0.5 hour after the dye injection, strong fluorescence was observed in cells most apical in the pallial VZ, which often overlapped with phospho-histone H3 (pH3, a mitosis marker) (Hendzel et al., 1997; Kim et al., 2017)-positive cells along the ventricular surface (Figure 1A). More than one third of the labeled cells were positive for pH3 (Figure 1F, -0.5 hour), although the mitosis rate should be underestimated by the time lag between dye injection and fixation. FT-labeled cells moved basally to leave the ventricular surface (Figures 1A-D). 3.5 hours after FT labeling, FT-labeled cells already left the ventricular surface and were almost never immunolabeled with pH3 (Figures 1B, F, -3.5 hours), suggesting that the labeling time window is less than several hours, compatible with previous observations (Telley et al., 2016). To visualize the difference of cellular population labeled with FT and thymidine analogues, we performed bolus injection of 5-ethynyl-2’-deoxyuridine (EdU, a thymidine analogue) into the intraperitoneal cavity of the mother mice at the same time with or 3-9 hours after FT labeling (Figures 1A-E). No cells were double-positive for FT labeling and EdU when EdU and FT labeling were performed simultaneously (Figures 1A, F, -0.5 hour). 9 hours after FT, some FT-labeled (FT+) cells were also labeled with EdU (EdU+), indicating that they reentered S-phase (Figures 1D, G, -9.5 hours). The slight differences of EdU+/FT+ between dorsomedial and dorsolateral cortices (Figure 1G, -9.5 hours, the definition of dorsomedial, dorsal and dorsolateral cortices, as well as cortical zones in this study are given in Figures S1A, B) might reflect differences in cell cycle length and/or proportion of direct neurogenesis (Polleux et al., 1997).

**Figure 1.**
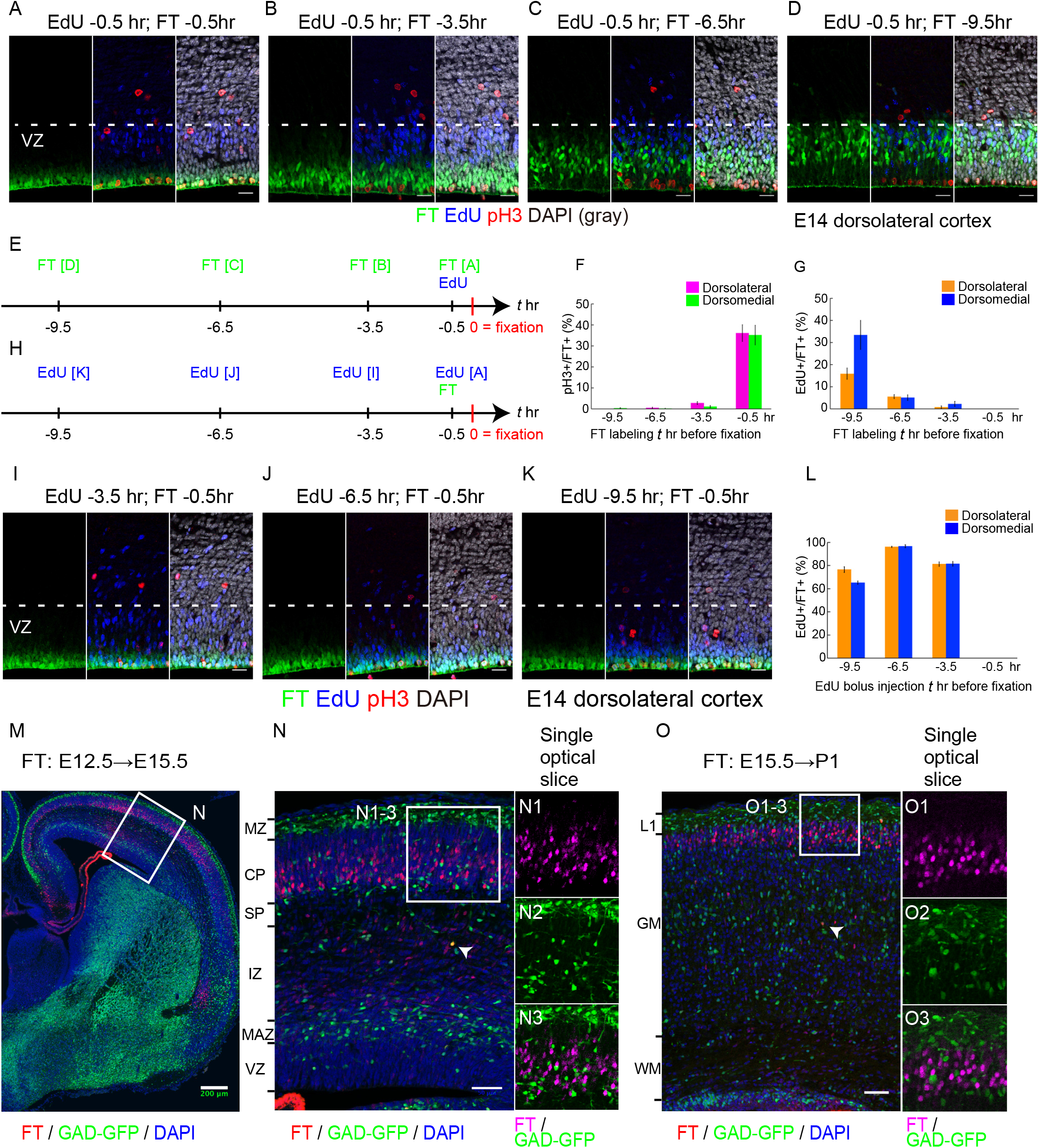
Characterization of cell population labeled with the FlashTag (FT) technology. **A-G:** 1 mM of 5- or 6-(N-Succinimidyloxycarbonyl) fluorescein 3’,6’-diacetate (CFSE) was injected into the lateral ventricles (LV) at embryonic day (E)14 ICR mice and fixed 0.5 (A), 3.5 (B), 6.5 (C), 9.5 (D) hours later. Intraperitoneal bolus injection of 5-Ethynyl-2’-deoxyuridine (EdU) was performed maternally 0.5 hours before fixation. Photomicrographs from the dorsolateral cortex were shown. In (A), FT-labeled cells positioned most apically and were often positive for phospho-histone H3 (pH3) (Figure 1F -0.5 hour, dorsolateral: 36.1± 4.0% [mean ± standard error of means], 339 cells from 5 brains; dorsomedial: 35.2 ± 4.7%, 249 cells from 5 brains) but negative for EdU administered at the same time (Figure 1G, -0.5 hour, dorsolateral: 0 ± 0%, 339 cells from 5 brains; dorsomedial: 0 ± 0%, 249 cells from 5 brains). The nuclei of EdU positive cells positioned basally in the VZ. 3.5 hours after FT injection, FT-labeled cells left the ventricular surface but still near it, and were no longer positive for pH3 (B) (Figure 1F, -3.5 hours, dorsolateral: 2.8 ± 0.7%, 530 cells from 5 brains; dorsomedial: 1.1 ± 0.5%, 415 cells from 5 brains). 6.5 hours after labeling, almost no cells were adjacent to the lateral ventricle (C). 9.5 hours after labeling, most of the labeled cells were in about basal two thirds in the VZ and double labeled for EdU, suggesting that some of them reentered the S-phase (D) (Figure 1G, -9.5 hours, dorsolateral: 15.9± 2.5%, 711 cells from 5 brains; dorsomedial: 33.4 ± 6.6 %, 546 cells from 5 brains). Schematic presentation of these experiments was shown in E. In (F), percentages of pH3+ cells out of the FT-labeled cells were shown. Magenta, pH3+ FT+/FT+ in the dorsolateral cortex. Green, pH3+ FT+/FT+ in the dorsomedial cortex. In (G), percentages of EdU+ cells out of FT-labeled cells were shown. Orange, EdU+ FT+/FT+ in the dorsolateral cortex. Blue, EdU+ FT+/FT+ in the dorsomedial cortex. **H-L:** EdU was administered 3 (I), 6 (J) and 9 (K) hours before FT labeling. 0.5 hours after FT labeling, brains were harvested. Schematic presentation of these experiments was shown in (H). Nuclei of the EdU labeled cells positioned more apically in brains in which EdU was administered 3.5 hours before fixation (I) compared with A, and some of the EdU labeled cells positioned at the ventricular surface to enter M phase (interkinetic nuclear migration). In brains which EdU was administered 6.5 (J) and 9.5 (K) hours before fixation, EdU-labeled cells positioned even more apically. In these mice treated with EdU 3-9 hours prior to FT, FT-labeled cells were often co-labelled with EdU (I-K) (Figure1L, dorsolateral, -9.5 hours: 76.6 ± 2.4%, 328 cells from 5 brains; -6.5 hours: 96.1 ± 0.5 %, 304 cells from 5 brains; -3.5 hours: 81.2 ± 1.9 %, 263 cells from 5 brains. Dorsomedial, -9.5 hours: 65.1 ± 1.5%, 369 cells from 5 brains; -6.5 hours: 96.7 ± 1.2 %, 217 cells from 5 brains; -3.5 hours: 81.5 ± 1.9 %, 287 cells from 5 brains). Note that EdU and FT never co-labeled when administered simultaneously (A). In the graph in (L), percentage of EdU+ cells out of FT-labeled cells were shown. Data for -0.5 hours in (L) corresponds to those for -0.5 hours in (G). Orange, EdU+ FT+/FT+ in the dorsolateral cortex. Blue, EdU+ FT+/FT+ in the dorsomedial cortex. **M-O:** CytoTell Blue was injected into the LV of the E12.5 (M-N) and 15.5 (O) GAD67-GFP brains. In E15.5 dorsolateral cortex labeled at E12.5, most of the labeled cells (red) were in the deep part of the cortical plate (CP) (M, N). Vast majority of the labeled cells were negative for GFP (E12.5-15.5 dorsolateral cortex, 93.3 ± 2.5%, 1653 cells from 3 brains) (N, N1-3). In postnatal day (P)1 dorsolateral cortex labeled at E15.5 (O), most of the labeled cells were found in the superficial gray matter. Again, vast majority of the labeled cells were negative for GFP (E15.5-P1, 95.5 ± 0.5 %, 1455 cells from 5 brains) (O, O1-3). Arrowheads in (N) and (O) show rare examples of cells positive for both FT and GFP. EdU, 5-Ethynyl-2’-deoxyuridine; DAPI, 4’,6-diamidino-2-phenylindole; FT, FlashTag; pH3, phospho-histone H3. GAD-GFP, Glutamate decarboxylase 67-green fluorescent protein, VZ, ventricular zone; MAZ, multipolar cell accumulation zone; IZ, intermediate zone; SP, subplate; CP, cortical plate; PCZ, primitive cortical zone; MZ, marginal zone; LI, cortical layer I; GM, gray matter; WM, white matter. Scale bars, 20 μm (A-D, I-K), 50 μm (N, O), 200 μm (M).

To further define the difference of labeled cellular population, we performed bolus injection of EdU into the intraperitoneal cavity of the mother mice at the same time with or 3-9 hours prior to FT (Figures 1H-L). Brains were harvested 0.5 hours after FT. Approximately 60 to 90% of FT-labeled cells in mice treated with EdU 3-9 hours prior to FT were co-labeled with EdU (Figure 1L, -3.5, -6.5, 9.5 hour). Since EdU is incorporated in the S-phase, these observations suggest that S-phase cells move apically to the ventricular surface by interkinetic nuclear migration over the course of several hours and are labeled with FT when they are around M-phase. Collectively, these observations suggest that FT labels cells around M phase on the ventricular surface almost specifically, and that FT can serve as a method to describe neuronal migration from the identical starting point, i.e., ventricular surface.

To investigate whether cortical interneurons were also labeled with FT, CytoTell Blue, another fluorescent dye used for FT labeling (Govindan et al., 2018; Telley et al., 2016), was injected into the LV of the E12.5 (Figures 1M, N) and E15.5 (Figure 1O) GAD67-GFP knock-in mouse brains (Tamamaki et al., 2003), in which interneurons are labeled with GFP. Examined several days after labeling, most of the FT-labeled cells were in the CP/gray matter and vast majority of them were negative for GFP. Many migrating cells in the lateral cortical stream (LCS) or “reservoir” (Bayer and Altman, 1991) were also mostly negative for GFP (Figures S1C, C1-C3). More ventrally, FT-labeled cells were identified in the caudal amygdaloid stream (CAS) (Remedios et al., 2007) and were negative for GFP (Figures S1C, C4-C6). GFP-labeled interneurons that migrated into the dorsal pallium were rarely labeled with FT, except for a small number of cells that were positive for both FT and GFP (Figures 1N, O, arrowheads), suggesting that most of the FT-labeled cells in the cortex are projection neurons when FT labeling is performed at E12.5-15.5.

Why did we observe only few FT-labeled interneurons in the cortex, although the ventral progenitors of the interneurons are also labeled with FT (Mayer et al., 2018) (Figures S1D, E)? We reasoned that frequent abventricular cell division might dilute the fluorescent dyes, losing fluorescent labeling. In fact, we performed immunohistochemistry against pH3 and confirmed that abventricular mitosis is very frequent in the ganglionic eminences (Figures S1D, E), compatible with previous reports (Katayama et al., 2013; Smart, 1976; Tan et al., 2016; Tan and Shi, 2013). We next reasoned that injection of a fluorescent dye into the SVZ might prevent loss of fluorescence by dilution upon abventricular mitosis. To address this, we injected CytoTell Blue into the parenchyma of the ganglionic eminences (the injection sites were retrospectively identified, e.g., the presumable injection site is shown by an asterisk in Figure S1F) in the *GAD67-GFP* mice at E12.5, and labeled interneuron progenitors far more strongly than intraventricular injections. FT and GFP-double labeled cells with tangential morphologies were distributed in the whole hemispheres, especially in the SVZ and marginal zone (MZ) of the dorsal pallium at E15.5 (Figures S1F, F1-10). These observations are compatible with the idea that FT-labeled interneuron progenitors in the VZ undergo mitosis in the SVZ, resulting in the loss of FT fluorescence in the migrating cortical interneurons when fluorescent dyes were injected intraventrically. Another group independently reported that the projection neurons occupied the majority of the FT-labeled cells using single-cell RNA-seq (Telley et al., 2019).

In summary, when fluorescent dyes were injected intraventricularly, the FT-labeled cells at early (E12.5) and late (E15.5) stages of neurogenesis were mostly non-GABAergic projection neurons. We do not preclude small subpopulation of interneurons from being labeled with FT, which will be described later in late and very late stages of neurogenesis (E15.5 and E17.0 cohort).

### FT visualizes clear regional differences in neuronal migration profiles in the cerebral cortex

During characterizing FT labeling, we noticed that there are regional differences in neuronal migration during development of the cerebral cortex using FT technology. We performed FT labeling at E14.5 and fixed two days later (Figure 2A). Many FT-labeled cells reached the top of the CP in the dorsomedial cortex while most of the labeled cells were still below the SP in the dorsolateral cortex (the lower border of the SP is shown by yellow dotted lines). This suggests that there are clear regional differences in neuronal migration profiles, e.g., times required for cells to reach the top of the CP, even where there is an underlying ventricular surface, when mitotic cells on the ventricular surface are selectively labeled with FT.

**Figure 2.**
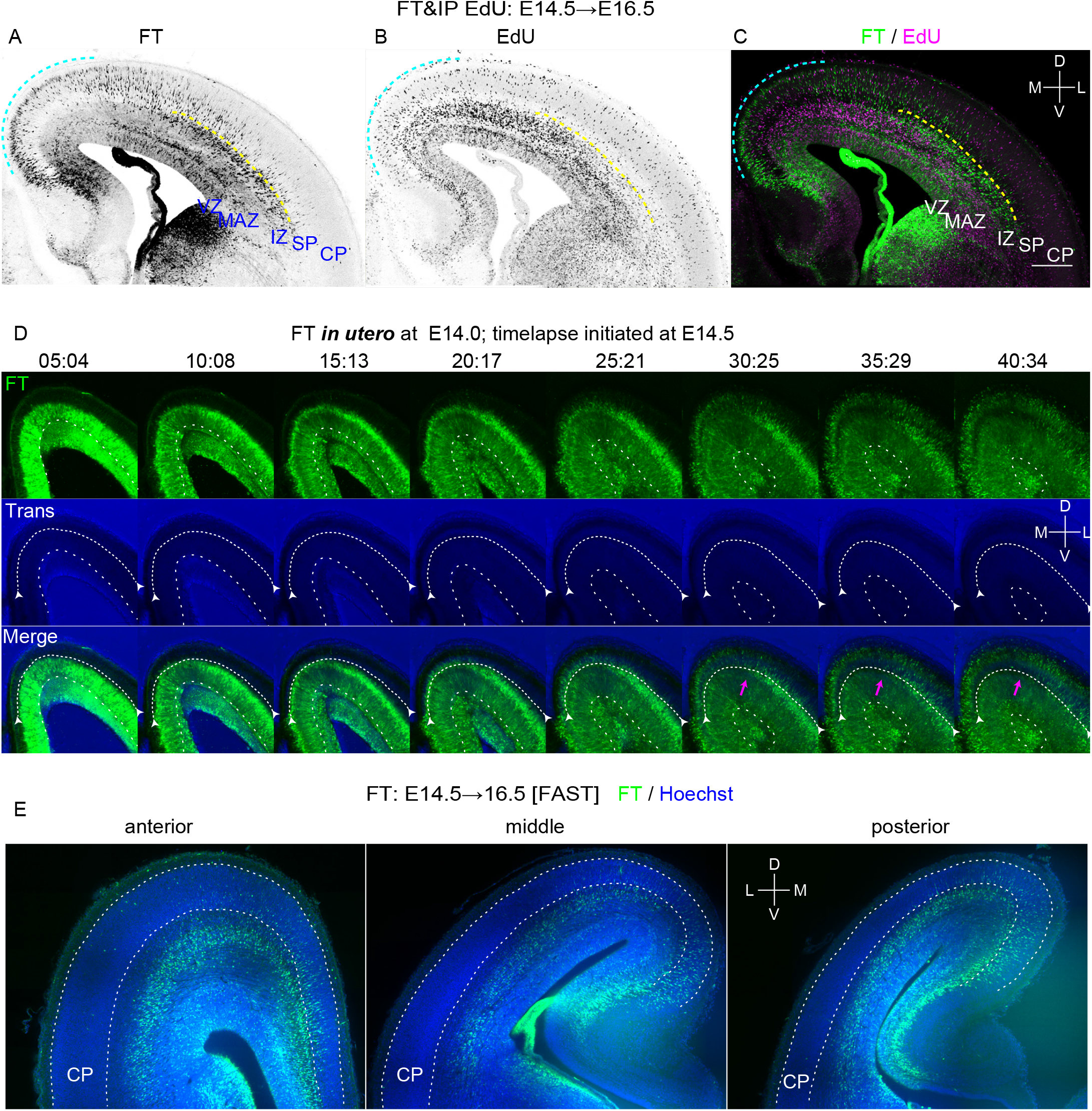
Regional differences in neuronal migration in the cerebral cortex revealed by FT. **A-C:** To visualize migration profile of the whole telencephalon, CFSE was injected into the ventricle of the E14.5 embryos and 5-Ethynyl-2’-deoxyuridine (EdU) was injected into the peritoneal cavity of the mother at the end of the surgery. Harvested at E16.5, many cells labeled with FT reached the superficial part of the CP in the dorsomedial cortex (cyan dotted line), while almost no cells reached the CP in the dorsolateral cortex (A, C). In the dorsolateral cortex, many neurons were just below the subplate (SP) (yellow dotted line). Such a clear difference in neuronal migration was not detected by EdU (B, C). **D:** FT labeling was performed at E14.0 and slice culture was prepared at E14.5. Labeled cells left the VZ and migrate in the MAZ in multipolar morphology (10:08-25:21). They gradually obtained polarity and migrate in the intermediate zone (20:17-30:25) and reached just below the SP (relatively dark band in the transmitted light channel, highlighted by white arrows). Neurons in the dorsomedial cortex (more medial than the magenta arrow) migrate smoothly to reach the most superficial part of the cortical plate(25:21-30:25), while in the dorsolateral cortex (more lateral than the magenta arrow), neurons seemed to sojourn transiently below the SP (clear in the regions lateral than the magenta arrow)(30:25-35:29). These cells subsequently migrated into the CP in the locomotion mode (35:29-40:34). **E:** FAST 3D imaging of E16.5 brains in which FT labeling was performed at E14.5. Anterior and posterior representative sections were shown in addition to a section at the interventricular foramen. Supplemental Movie 1 shows a whole 3D movie taken from this brain. EdU, 5-Ethynyl-2’-deoxyuridine; DAPI, 4’,6-diamidino-2-phenylindole; FT, FlashTag; IP, intraperitoneal injection; VZ, ventricular zone; MAZ, multipolar cell accumulation zone; IZ, intermediate zone; SP, subplate; CP, cortical plate; M, medial; L, lateral; D, dorsal; V, ventral. Scale bars, 200 μm.

To compare migration profiles visualized with FT labeling with those with thymidine analogues, EdU was also administered at the same time with FT labeling. The distribution of EdU-positive cells was similar to that reported previously (Bayer and Altman, 1991) as expected, and the distribution of labeled cells did not clearly differ between the dorsomedial and dorsolateral region when examined where there is an underlying ventricular zone (Figures 2B, C). The EdU-labeled neurons in the dorsolateral CP (Figure 2B) should have passed the SP to enter the CP earlier than FT-labeled cells (Figure 2A), although M-phase labeled (FT-labeled) cells should have started migration earlier than S-phase labeled (EdU-labeled) cells if they were labeled in the VZ. Therefore, EdU-labeled neurons in the dorsolateral CP in Figure 2B would have been in the S-phase in the SVZ at the time of FT labeling and EdU administration. We previously reported that mitotically-active population leaving the VZ (rapidly exiting population, or REP) (Tabata et al., 2009), most of which corresponds to the basal progenitors and glial progenitors (Tabata et al., 2009; Tabata et al., 2012), are abundant in the dorsolateral cortex than dorsomedial cortex. This population would have contributed to the EdU-labeled cells in the dorsolateral CP.

To further characterize these regional differences, we performed time-lapse imaging of the FT-labeled cells (Figure 2D). In the dorsolateral cortex, labeled cells left the VZ to enter and accumulate in the multipolar cell accumulation zone (MAZ), a zone enriched in postmitotic multipolar cells (Tabata et al., 2009; Tabata et al., 2012)(Figure 2D, 10:08-25:21). They then migrated through the IZ and transiently sojourned just below the SP (Figure 2D, 30:25-35:29) before entering the CP (Figure 2D, 40:34). This sojourning behavior below the SP would correspond to the stationary period (Ohtaka-Maruyama et al., 2018). To note, this sojourning behavior was not clear in the dorsomedial cortex. To visualize the migratory profile in 3 dimensions (3D), we injected the dye at E14.5 and fixed about two days later, and the brains were subjected to 3D FAST imaging (Seiriki et al., 2017; Seiriki et al., 2019). The mediolateral difference in the migratory profile was preserved along the anterior-posterior axis (Figure 2E, Supplemental movie. 1). The difference was somewhat clearer in posterior cortex (presumptive retrosplenial - visual cortex) than anterior (presumptive medial prefrontal cortex – somatosensory cortex). These observations suggest that this mediolateral difference in neuronal migration profiles may, at least in part, result from transient pause just below the dorsolateral SP.

### Regional migratory/positional profiles differ from neuronal birthdates at the ventricular surface

It was previously reported that early and late-born neurons migrate differently (Hatanaka et al., 2004). Do both early-born neurons and late-born neurons show similar regional differences? Or do regional differences have birthdate-dependent characters? To better understand migration profiles of neurons born at different embryonic stages, we injected CFSE at E10.5, 11.5, 12.5 13.5, 14.5, 15.5, 17.0 and fixed chronologically. Subsequent observation was carried out on the coronal section in which the interventricular foramen was visible, respectively.

### E10.5 cohort

One day after injection, at E11.5, some labeled cells were already located in the PP in both dorsomedial and dorsolateral cortex (Figures 3D, E, Figure S2A), although the formed PP is very thin especially in the dorsomedial cortex (Figure 3D). Many other labeled cells were still in the VZ, which consists of densely-packed, radially oriented (Boulder-Committee, 1970) Pax6-positive (Englund et al., 2005) nuclei of radial glia. Two days after injection, at E12.5, most of the labeled cells were in the PP (Figure 3A, D, E, Figure S2B). Mediolateral migratory differences were not clear in these observations.

**Figure 3.**
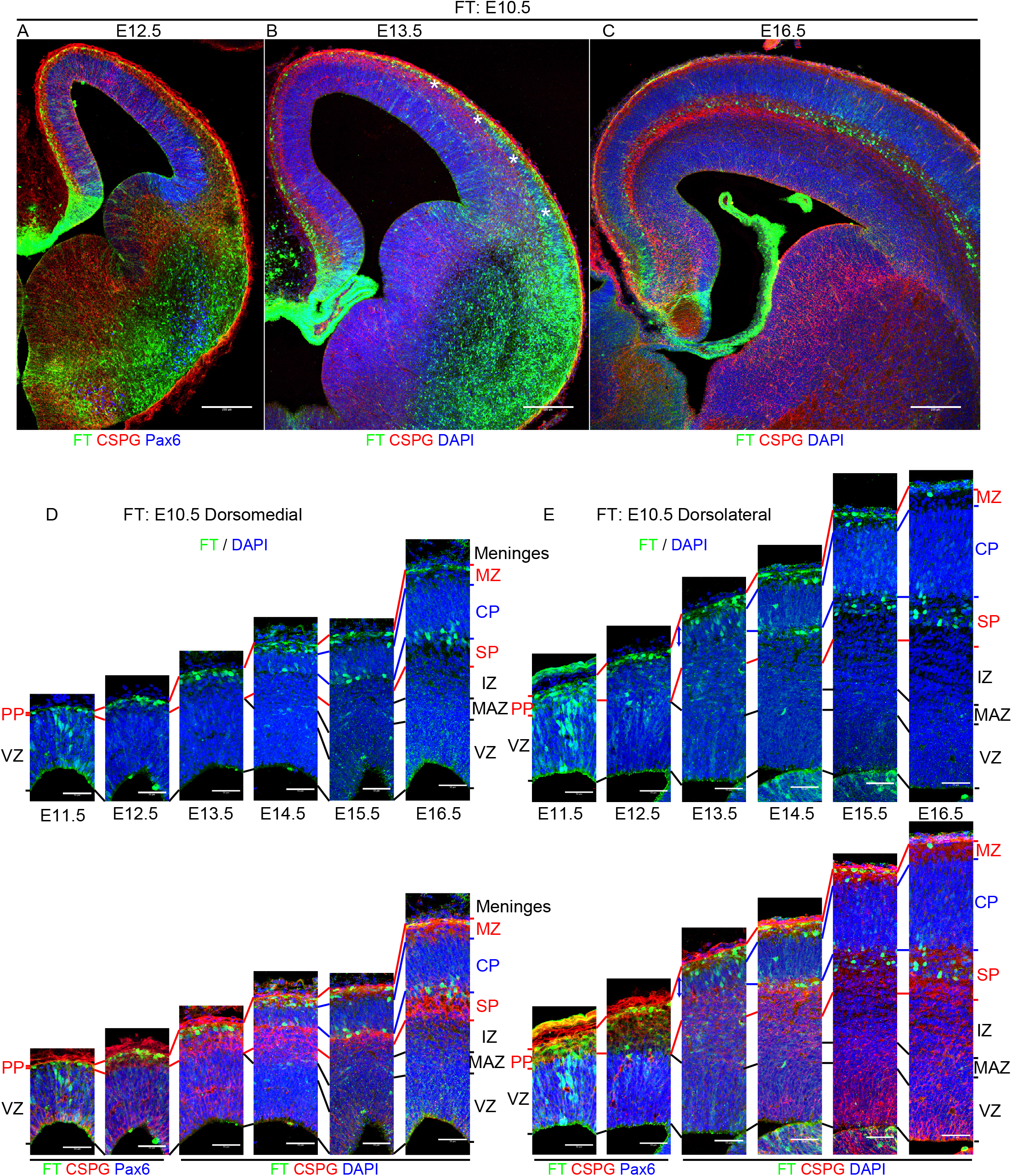
Cohort of cells born at E10.5. **A-E:** Coronal sections of 12.5 (A), 13.5 (B) and 16.5 (C) brains labeled at E10.5. See also Figure S2 for coronal section from E11.5 to 16.5 shown with FT and DAPI. Higher magnification pictures from the dorsomedial cortex and dorsolateral cortex from E11.5 to 16.5 were shown in (D) and (E), respectively. As early as E11.5, some cells were found in the preplate (PP), which was very thin in the dorsomedial cortex, as well as in the VZ (D, E, Figure S2A). At E12.5, many cells were in the PP, sometimes in a tangential morphology (A, D, E). At E13.5, the CSPG and nuclear staining showed PP splitting proceeded in a lateral-to-medial direction and the CP (asterisks) was observed in the dorsolateral cortex but not in the dorsomedial cortex (B). In the dorsomedial cortex, labeled cells were in the PP, often in somewhat round morphology (D). In the dorsolateral cortex, on the other hand, many labeled cells were located in the CP (shown with blue arrows) and MZ (E). Note that few cells were found below the CP identified by nuclear and CSPG staining (B, E, Figure S2C). At E14.5, thin CP was identified in the dorsomedial cortex as well (D, E, Figure S2D). Some labeled cells were in the deep part of the CP in the dorsomedial cortex, but many labeled cells were still in the MZ (D). In the dorsolateral cortex, many labeled cells were found near the boundary between the CP and SP (D, Figure S2D). At E15.5, labeled cells were found at the boundary between SP and CP as well as MZ in the dorsomedial cortex (D, Figure S2E, S2E’), which is similar to the dorsolateral cortex of the E14.5 (E, Figure S2D). In the E15.5 dorsolateral cortex, many labeled cells were in the CSPG-positive SP (D, E, Figure S2E, S2E’). At E16.5, in both the dorsomedial and dorsolateral cortex, labeled cells were mainly found in the SP (C, D, E). Some cells were also found in the MZ (D, E, Figure S2G). Note that the CSPG staining in the SP showed some double-track immunoreactivity strong just above and below the distinct cell layer in the SP in dorsal and dorsolateral cortex at E15.5-E16.5 (E). Emergence of the labeled cells in the SP seems to coincide with this emergence of distinct layer. DAPI, 4’,6-diamidino-2-phenylindole; FT, FlashTag; VZ, ventricular zone; PP, preplate; MAZ, multipolar cell accumulation zone; IZ, intermediate zone; SP, subplate; CP, cortical plate; MZ, marginal zone. Scale bars, 200 μm (A-C) and 50 μm (D, E).

Three days after injection, at E13.5, the CSPG (Bicknese et al., 1994) and nuclear staining showed PP splitting proceeded in a lateral-to-medial direction and an emergence of the cortical plate was apparently recognized in the dorsolateral cortex (asterisks in Figure 3B, a blue arrow in Figure 3E) but not in the dorsomedial cortex (Figures 3B, D)(upper and lower borders of the formed CP are shown by blue lines in Figures 3D and 3E), which is consistent with the lateral-to-medial neurogenic gradient. In the dorsolateral cortex, most of the labeled cells were in the CP and MZ, whereas in the dorsomedial cortex, most of the labeled cells were in the PP. Note that strongly-labeled cells were hardly found in the SP just below the CP at E13.5 (Figures 3B, D, E, Figure S2C). At E14.5-15.5 in the dorsomedial cortex, labeled cells were found at the boundary between SP and CP as well as in the MZ (Figures 3D, Figures S2D, S2E), which was similar to the dorsolateral cortex of the E14.5 (Figures 3E, Figure S2D). In the E15.5 dorsolateral cortex, many labeled cells were distributed in the CSPG-positive SP below the CP (Fig. 3E, Figure S2E). At E16.5, in both the dorsomedial and dorsolateral cortex, labeled cells were mainly found in the SP (Figures 3C, D, E, Figure S2F) and were Tbr1 (Hevner et al., 2001)-positive (Figure S2G), suggesting that they are of pallial origin. Some cells were also found in the MZ (Figures 3C, D, E, Figure S2G) and were also positive for Reelin (Ogawa et al., 1995) (Figures S2G), suggesting that FT labeling at E10.5 mainly labels Tbr1-positive SP cells and Cajal-Retzius cells. These observations suggest that at least some future SP neurons in the PP are located in the CP and MZ when the CP begins to be formed. They might eventually move down to the SP layer in a lateral-to-medial fashion. This view is compatible with previous observations (Bayer and Altman, 1991; Osheroff and Hatten, 2009; Saito et al., 2019). Recent observations have shown that future SP neurons migrate tangentially in the PP (Pedraza et al., 2014; Saito et al., 2019), but FT failed to explicitly detect tangential migration of the future SP neurons probably because FT labels the whole hemispheres.

In summary, E10.5 cohort reached the PP in less than one day after they exited the VZ in both dorsomedial and dorsolateral cortices. Among the E10.5 cohort, future SP neurons formed a distinct layer below the CP in a lateral-to-medial fashion, reflecting the well-described neurogenic gradient.

### E11.5 cohort

As early as half a day after injection, at E12.0, most of the labeled cells were located in the VZ, and some cells were in the CSPG-positive PP in both the dorsomedial (Figures 4D, E, an arrowhead in Figure 4D, Figure S3A) and dorsolateral cortex (Figures 4D, E, arrowheads in Figure 4E, Figure S3A). At E12.5 and 13.0, more labeled cells were found in the PP in both dorsomedial (Figures 4A, B, D, Figures S3B, S3C) and dorsolateral cortex (Figures 4A, B, E, Figures S3B, S3C) as well as in VZ. Many labeled cells reached just beneath the meninges. At E 13.5, in the dorsomedial cortex, where PP splitting has not occurred yet at this stage, many neurons were located in the PP just beneath the meninges (Figure 4D, Figure S3D). However, in the dorsolateral cortex, where the PP was split, many labeled cells showed radial (parallel to the apicobasal axis) alignment in the newly formed CP (Figure 4E, Figure S3D). The formation of the CP coincided with this radial alignment of the labeled cells at E13.5 in the dorsolateral cortex and E14.5 in the dorsomedial cortex (Figures 4D, E, Figure S3E). At E15.5, some cells were located in the MZ, others stayed in the deep part of the CP and expressed a deep layer marker Ctip2 (Arlotta et al., 2005), and still others emerged below the CP (Figures 4C, D, E, Figures S3F, S3G, S3H).

**Figure 4.**
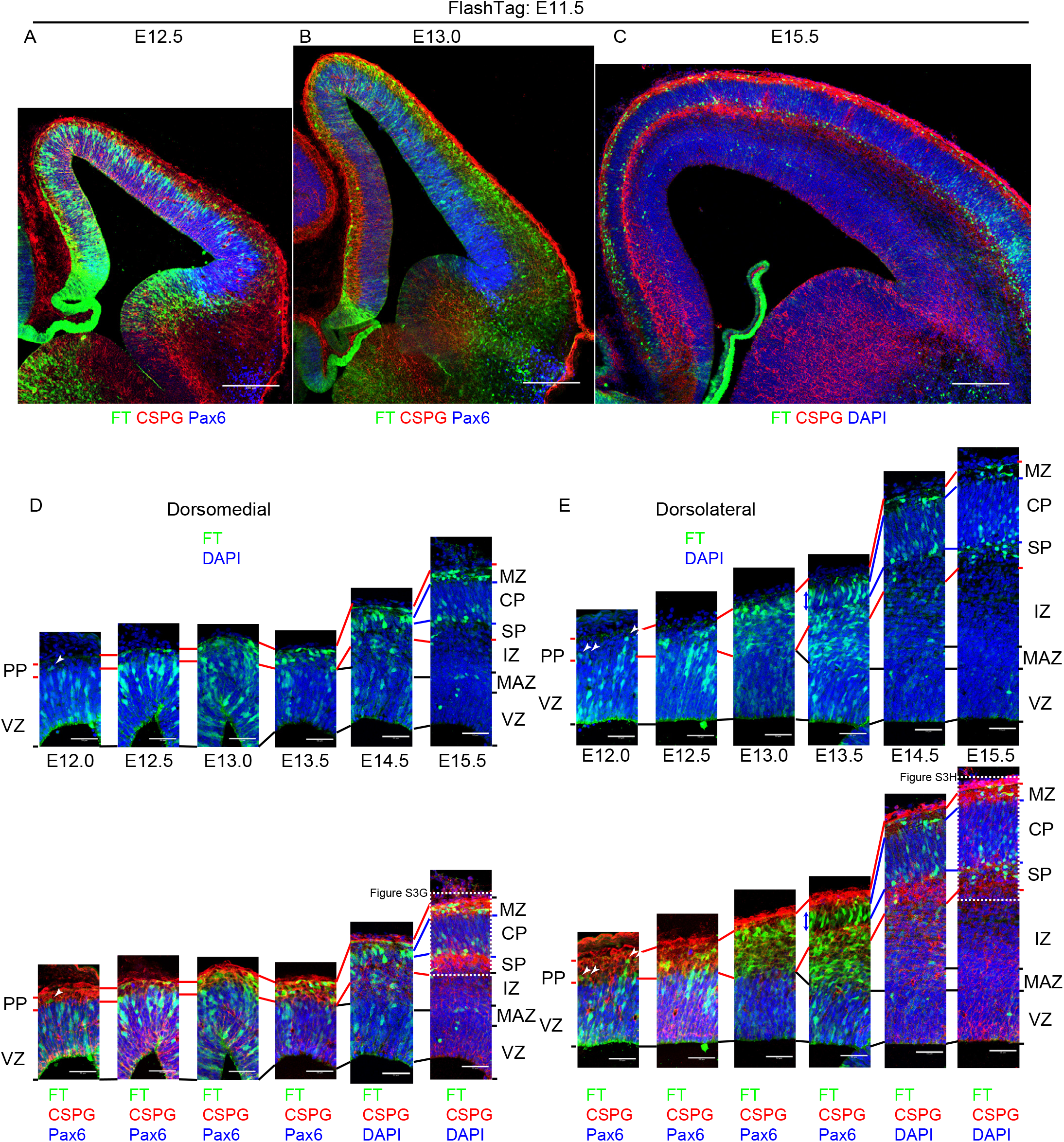
Cohort of cells born at E11.5. **A-E:** Coronal sections of E12.5 (A), 13.0 (B) and 15.5 (C) brains labeled at E11.5. Higher magnification pictures from the dorsomedial cortex and dorsolateral cortex of E12.5 through E15.5 were shown in (D) and (E), respectively. See also Figure 3S for low magnification pictures of brains fixed at E12.5 through E15.5. At E12.0, most of the labeled cells were located in the VZ, and some cells were in the CSPG-positive PP in both the dorsomedial and dorsolateral cortex (D, E, arrowheads, Figure S3A). At E12.5 and 13.0, more labeled cells were found in the PP in both dorsomedial (A, B, D, Figures S3B, S3C) and dorsolateral cortex (A, B, E, Figures S3B, S3C). At E 13.5, in the dorsomedial cortex, where PP splitting does not occur yet at this stage, many neurons reached the PP just beneath the meninges (D, Figure S3D). Many labeled cells were located in the newly formed CP and intermediate zone (IZ) in the dorsolateral cortex (E, Figure S3D). At E14.5, many cells were in the newly formed CP in both dorsomedial and dorsolateral cortex (D, E, Figure S3E). At E15.5, many cells were in the lower part of CP and, to lesser extent, MZ (C, D, E, Figures S3F, S3G, S3H). Some cells were also found in the SP in the dorsolateral cortex (C, E, Figures S3F, S3H). DAPI, 4’,6-diamidino-2-phenylindole; FT, FlashTag; VZ, ventricular zone; PP, preplate; MAZ, multipolar cell accumulation zone; IZ, intermediate zone; SP, subplate; CP, cortical plate; MZ, marginal zone. Scale bars, 200 μm (A-C) and 50 μm (D, E).

In summary, E11.5 cohort reached the PP soon after they exited the VZ in both dorsomedial and dorsolateral cortices, like E10.5 cohort. The formation of radial alignment occurred after they reached just below the meningeal surface in a lateral-to-medial fashion, in parallel with the formation of the CP. As in the E10.5 cohort, future SP neurons formed a distinct layer below the CP after the CP formed in the E11.5 cohort.

### E12.5 cohort

As early as half a day after injection, at E13.0, many labeled cells were in the VZ, but a small number of labeled cells were also found in the PP in the dorsomedial cortex (Figures 5A, G, Figure S4A, S4G). The latter cells were often weakly positive for Pax6 (Figure S4G, arrowheads) (number of FT+/Pax6+ cells, 9.0 ± 2.0 cells [mean ± standard error of means] /dorsomedial low power field, *n* = 4 brains). On the other hand, in the dorsolateral cortex, many labeled cells were located in zones just above the VZ in addition to the VZ, and they are mostly negative for Pax6 (Figures 5A, H, Figure S4A, S4H; arrows). FT+ / Pax6+ cells outside of the VZ were relatively rare in the dorsolateral cortex (number of FT+/Pax6+ cells, 1.6 ± 0.8 cells/dorsolateral low power field, *n* = 4 brains). One day after injection, at E13.5, more labeled cells were in the PP in addition to the VZ in the dorsomedial cortex and labeled cells in the PP were no longer positive for Pax6 (Figures 5B, G, Figure S4B). In the dorsolateral cortex, the incipient CP appears at this stage (E13.5) (See also Figure 3B), and majority of the labeled cells were below the CP and still migrating in the IZ (Figures 5B, H, Figure S4B). Some of them entered and were radially aligned in the CP; others were migrating in the IZ at E14.0 (Figures 5C, H, Figure S4C). In the dorsomedial cortex of E14.0, however, when the incipient CP is about to be formed, many labeled cells had already reached just beneath the meningeal surface (Figures 5C, G, Figure S4C). The radial alignment of the labeled cells coincided with the formation of the CP at E14.5 in the dorsomedial cortex (2 days after injection) (Figures 5D, G, Figure S4D), as in the E11.5 cohort. Most of the labeled cells occupy the CP at E15.5 at both dorsomedial and dorsolateral cortex (Figures 5E, G, H, Figure S4E), and some emerged in the SP at E16.5 in the dorsomedial cortex (Figures 5F, G, Figure S4F).

**Figure 5.**
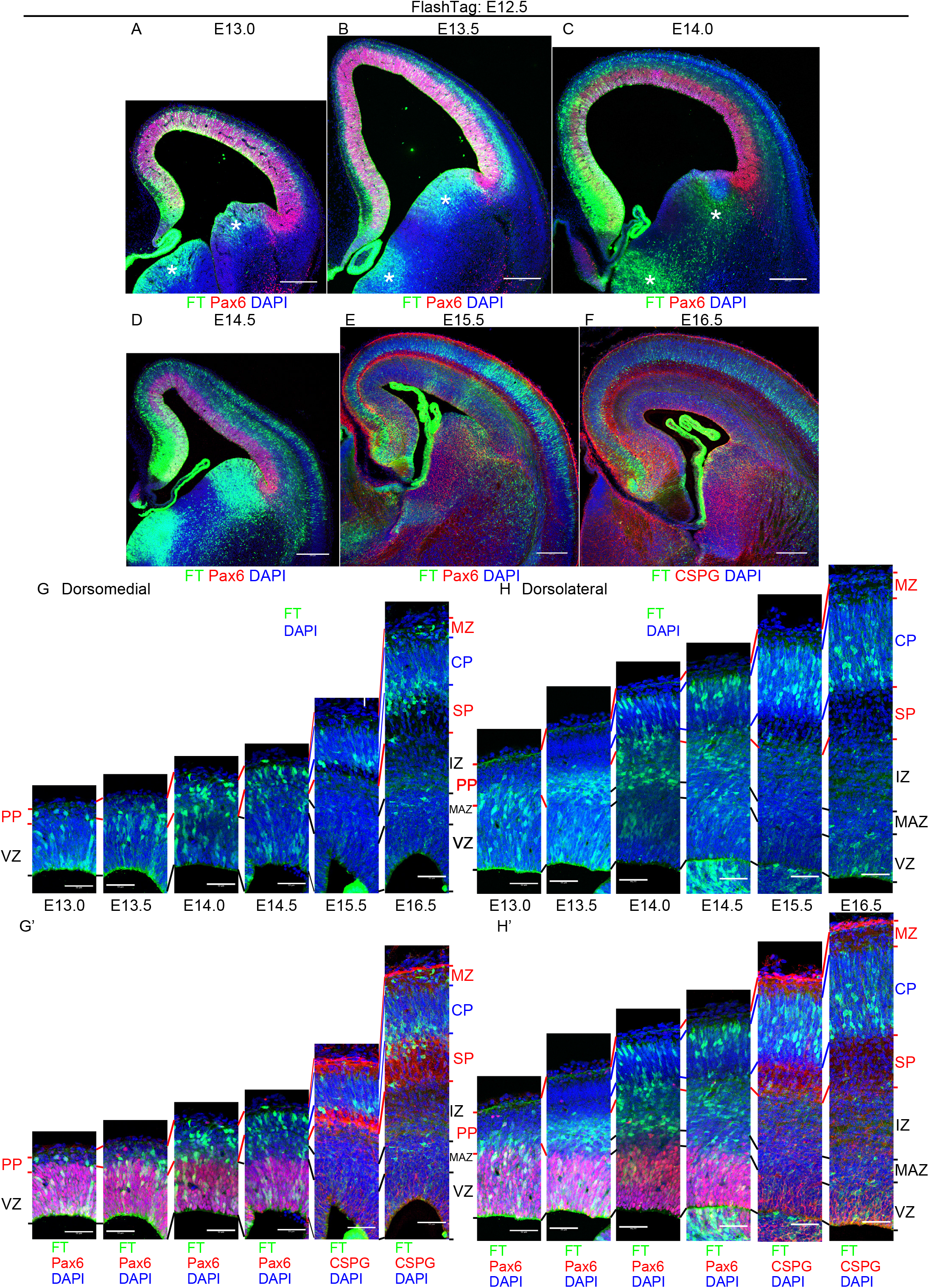
Cohort of cells born at E12.5. **A-H:** Coronal sections of E13.0 (A), 13.5 (B), 14.0 (C), 14.5 (D), 15.5 (E), and 16.5 (F) brains labeled at E12.5. Higher magnification pictures from the dorsomedial cortex and dorsolateral cortex were shown in (G) and (H), respectively. In the dorsomedial cortex at E13.0, many labeled cells were in the VZ, but a small number of labeled cells were also found in the PP (A, G, Figure S4G). At E13.5, more labeled cells were in the PP in addition to the VZ in the dorsomedial cortex (B, G). At this stage, the incipient CP appears in the dorsolateral cortex, and many labeled neurons were migrating in the IZ (B, H). In the dorsomedial cortex of E14.0, when the incipient CP is beginning to be formed, some labeled cells reached just beneath the meningeal surface, while others seemed to be still migrating (C, G). In the dorsolateral cortex, too, many neurons reached the superficial part of the CP, while others were still migrating in the IZ and CP (C, H). At E14.5, labeled cells in the dorsomedial cortex began to be oriented radially just beneath the MZ (D, G). In the dorsolateral cortex, many strongly labeled cells were located in the cortical plate in addition to the IZ (D, H). At E 15.5, most of the labeled cells distributed not only the superficial CP, but also in the deep part of the CP in both the dorsomedial and dorsolateral CP, suggesting that some of them began to move deeper (E, G, H). At E16.5, the main population of the labeled cells was located in the somewhat deeper part of the CP in both the dorsomedial and dorsolateral cortex. In the dorsomedial cortex, many labeled cells were also distributed in the SP. DAPI, 4’,6-diamidino-2-phenylindole; FT, FlashTag; VZ, ventricular zone; PP, preplate; MAZ, multipolar cell accumulation zone; IZ, intermediate zone; SP, subplate; CP, cortical plate; MZ, marginal zone. Scale bars, 200 μm (A-F), 50 μm (G, H).

Taken together, we observed slight signs of mediolateral differences in migration profiles of neurons labeled at E12.5, that is, dorsomedial E12.5 cohort reached the outermost region of the PP just beneath the pial surface relatively soon after they left the VZ, while dorsolateral E12.5 cohort slowly migrate in the lower part of the PP or IZ before they enter the CP. Radial alignment of the labeled cells, on the other hand, occurred in a lateral-to-medial fashion in parallel with the formation of the CP.

### E13.5 cohort

Half a day after injection, at E14.0, most of the labeled cells were located in or just above the VZ, i.e. the multipolar cell accumulation zone (MAZ), a zone enriched in postmitotic multipolar cells (Tabata et al., 2009; Tabata et al., 2012), in both dorsomedial and dorsolateral cortex (Figures 6A, F, G, Figure S5A). One day after injection, at E14.5, many labeled neurons were migrating in the IZ below the CSPG-positive SP (Figures 6B, F, G, Figure S5B). One and a half days after injection, at E15.0, many labeled cells reached the top of the CP in the dorsomedial cortex (Figures 6C, F, Figure S5C), while in the dorsolateral cortex, few cells reached the CP and many cells were still migrating in the superficial IZ or beneath the SP (Figures 6C, G, Figure S5C). In the dorsolateral cortex, it was at E15.5-16.5 when most of the labeled cells reached the superficial CP (Figures 6D, E, G, Figure S5D, S5E). There observations suggest that there are clear regional differences in times required for neurons to reach the CP for the E13.5 cohort. At E16.5, E17.5 and E18.5, labeled neurons were overtaken by neurons presumptively born later to settle in the deep part of the CP (Figures 6E, F, G, Figures S5F, G) in dorsomedial and dorsolateral cortices. In the ventrolateral cortex, some labeled neurons were still in the reservoir and others were migrating out the reservoir to the insular and piriform CP (Figures S5F, F1) at E17.5, compatible with the previous observation that neurons which migrate along the lateral cortical stream take longer to reach their final destinations (Bayer and Altman, 1991). Labeled neurons were also observed in the presumptive CAS (Remedios et al., 2007) (Figures 5F2).

**Figure 6.**
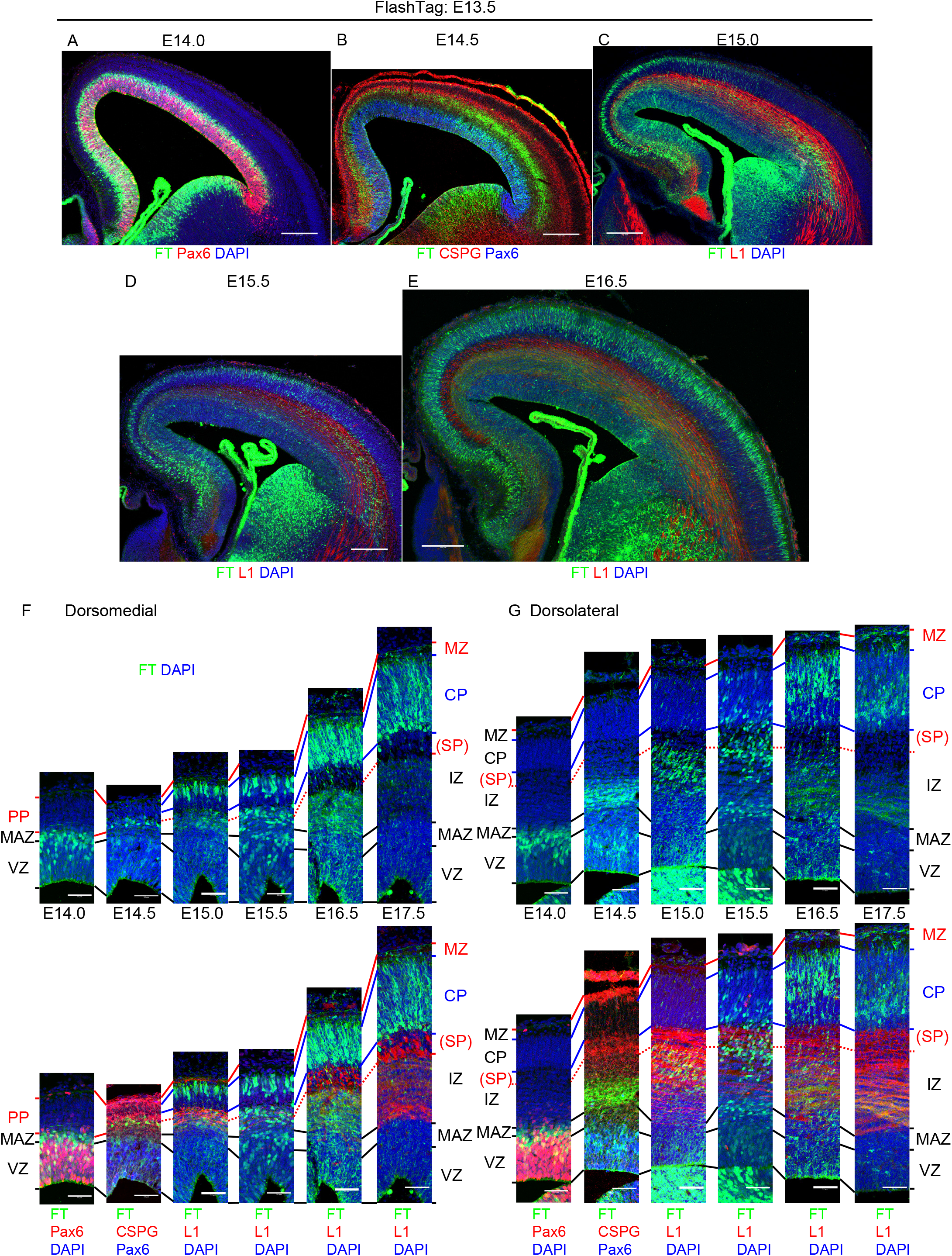
Cohort of cells born at E13.5. **A-G:** Coronal sections of E14.0 (A), 14.5 (B), 15.0 (C), 15.5 (D) and 16.5 (E) brains labeled at E13.5. Higher magnification pictures from the dorsomedial cortex and dorsolateral cortex from E14.0-17.5 were shown in (F) and (G), respectively. Lower magnification pictures of E17.5 and E18.5 are shown in Figure S5. Coronal sections of FT-labeled brains of E14.0 through E18.5 stained with nuclear staining were also shown in Figure S5. At E14.0, most of the labeled cells were located in the VZ and zones just above the VZ in both dorsomedial and dorsolateral cortex (A, F, G). At E14.5, many labeled neurons were migrating in the IZ below the SP as revealed by immunohistochemistry for CSPG (B, F, G). At E15.0, majority of the labeled cells reached just beneath the pial surface in the dorsomedial cortex (C, F) but most of the labeled cells in the dorsolateral cortex were in the IZ below the SP (C, G). At E15.5, some of the labeled cells entered the CP while many neurons were still migrating in the IZ and SP in the dorsolateral cortex (D, G). At E16.5, most of the labeled cells in the dorsomedial cortex were located in the CP (E, F). Most of the labeled cells in the dorsolateral cortex reached the superficial part of the CP. Note that FT-labeled axon bundles were in the IZ. At E17.5, many strongly labeled neurons located in the deeper part of the CP, suggesting that later-born neurons passed through the neuronal layers that were born at E13.5 (F, G, Figure S5F, S5G). DAPI, 4’,6-diamidino-2-phenylindole; FT, FlashTag; VZ, ventricular zone; PP, preplate; MAZ, multipolar cell accumulation zone; IZ, intermediate zone; SP, subplate; CP, cortical plate; MZ, marginal zone. Scale bars, 200 μm (A-E), 50 μm (F, G).

In summary, cells labeled at E13.5 reached just beneath the meningeal surface in about 1.5 days in the dorsomedial cortex, while those in the dorsolateral cortex took longer to enter the CP and thus to reach just beneath the meningeal surface. This areal difference was similar to that observed in E14.5 cohort in Figure 2, and likely to be explained by transient sojourning below the SP, at least in part.

### E14.5 cohort

To confirm if regional migratory differences observed in the slice time-lapse imaging are indeed observed *in vivo*, and to characterize regional differences better histologically, we performed FT labeling at E14.5 and fixed chronologically (Figures 7A-I, Figures S6A-H). Half a day after injection, at E15.0, most of the labeled cells were in the VZ (Figure 7A, H, I, Figures S6A). Small number of labeled cells left the VZ mainly in the dorsolateral cortex (Figure 7H, J) (dorsomedial, 3.7 ± 0.9 cells/low power field, *n* = 3 brains; dorsolateral, 23.0 ± 3.5 cells/low power field, *n* = 3 brains). They have a long ascending process and a retraction bulb, and mitotically active as shown by Ki-67 immunoreactivity (Figure 7J), which presumably corresponds to the mitotically-active rapidly exiting population (REP) that we reported previously (Tabata et al., 2009). They were also positive for stem cell markers Pax6 and Sox2, although they were outside of the VZ (Figure 7J) (Pax6 positive, dorsomedial, 91.7 ± 8.3%, 11 cells from 3 brains; dorsolateral, 80.8 ± 2.9%, 69 cells from 3 brains), suggesting that most of the cells in this population has a feature of mouse outer radial glial cells (moRG) (Shitamukai et al., 2011; Vaid et al., 2018; Wang et al., 2011). The progeny of this population was often difficult to identify, probably because the fluorescent signals decrease upon mitosis in the SVZ. One day after injection, at E15.5, the major population of the labeled cells left the VZ and accumulated in the MAZ (Figures 7B, H, I, Figures S6B). One and a half days after injection, at E16.0, most of the labeled cells were migrating in the IZ (Figures 7C, H, I, Figures S6C). Until this timepoint, mediolateral migratory difference of the major population was not so clear. Two days after injection, at E16.5, however, many cells reached the most superficial part of the CP in the dorsomedial cortex, while the majority were still migrating in the IZ just beneath the SP in the dorsolateral cortex (Figures 7D, H, I, Figures S6D), showing a clear mediolateral difference in migration. These observations are consistent with the view that mediolateral migratory difference is attributable to the migratory behavior of neurons in the dorsolateral IZ or just beneath the dorsolateral SP. In the dorsomedial cortex, most of the labeled cells settled in the most superficial CP, or PCZ, a zone composed of densely packed immature neurons (Sekine et al., 2011; Sekine et al., 2012; Shin et al., 2019), at E16.5-17.5 (Figures 7D, E, H). By E18.5, they were overtaken by presumptive later-born neurons and positioned in a slightly deeper part of the CP as NeuN-positive mature neurons in the dorsomedial cortex (Figures 7F, H). In the dorsolateral cortex, it took one more day to reach the CP (Figures 7E, I, Figures S6E) and settled in the PCZ (double headed arrows in Figures 7G, H, I) at E17.5-18.5 (Figures 7E, F, I, Figures S6E, S6F). At P0.5, they positioned in a slightly deeper part of the CP as NeuN-positive mature neurons (Figure 7G) by being taken over by immature neurons in the PCZ. At P7, labeled neurons mainly positioned in layer II/III in the dorsomedial and lateral cortices and in layer IV in the dorsolateral cortex (Figure S6H).

**Figure 7.**
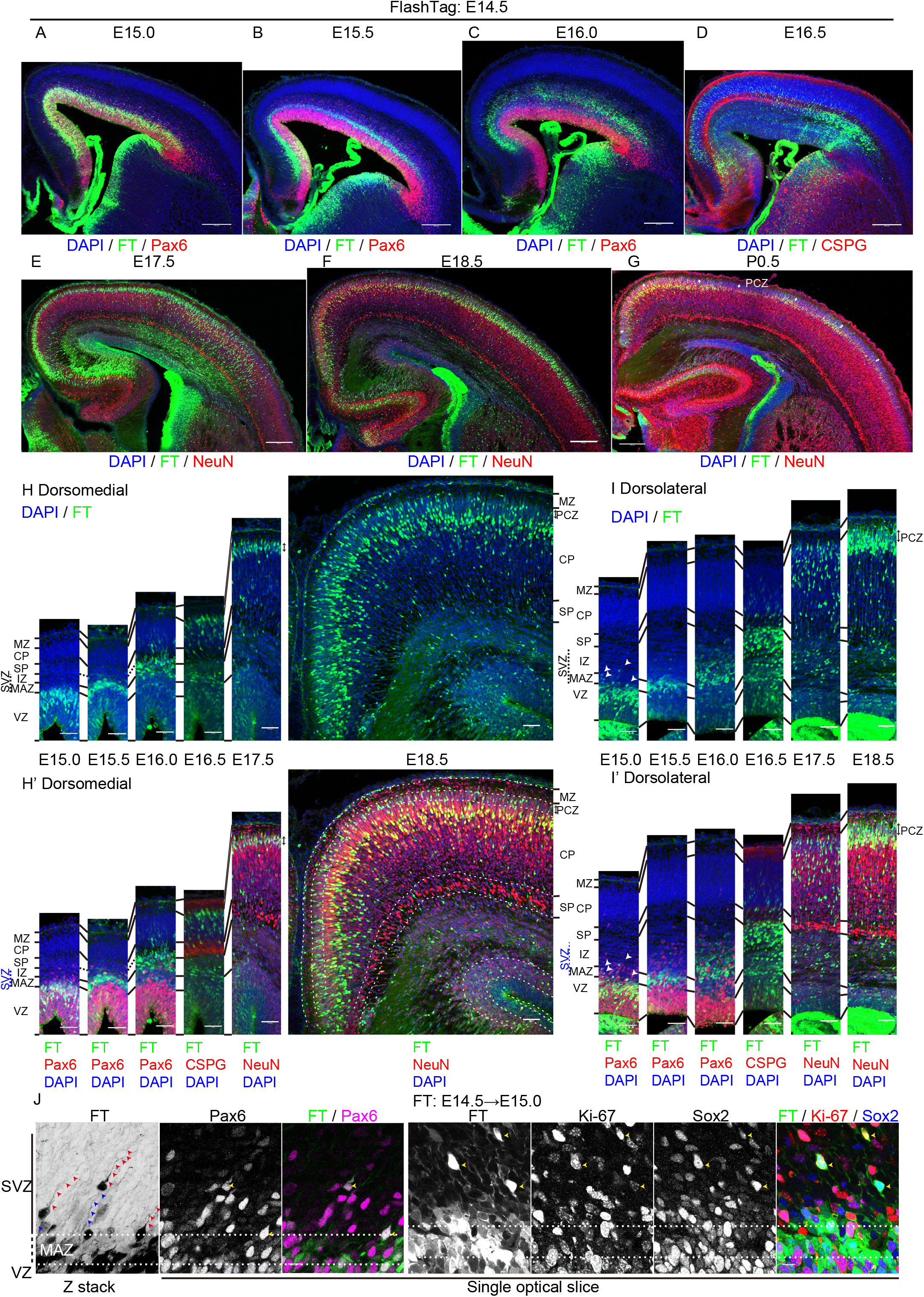
Cohort of cells born at E14.5. **A-J:** Coronal sections of E15.0 (A), 15.5 (B), 16.0 (C), 16.5 (D), 17.5 (E), 18.5 (F) and P0.5 (G) brains labeled at E14.5. Higher magnification pictures from the dorsomedial cortex and dorsolateral cortex were shown in (H) and (I), respectively. Higher magnification of the apical part of the dorsolateral cortical wall of E15.0 (0.5 days after injection) brains was shown in (J). At E15.0, most of the labeled cells were located in the VZ in both dorsomedial and dorsolateral cortex (A, H, I). Some labeled cells were located outside of the VZ in the dorsolateral cortex (A, I, J) but such cells were not frequently found in the dorsomedial cortex (A, H). The labeled cells that were located basally often had a long ascending process (red arrowheads, J, left) as well as some retraction bulb (blue arrowheads) and were immunoreactive for Pax6, Sox2 and Ki-67 (yellow arrowheads, J, right). Note that the ascending processes were so long that it was difficult to observe full length in the IZ crowded with radial fibers, which are also labeled with FT. At E15.5, majority of the labeled neurons were located in the MAZ in the multipolar morphology in both the dorsomedial and dorsolateral cortex (B, H, I). At E16.0, most of the labeled cells were in the IZ (C, H, I). At E16.5 in the dorsomedial cortex, many cells reached the most superficial part of the CP (D, H). In the dorsolateral CP, on the other hand, most of the labeled cells were migrating in the IZ just beneath the SP (D, I; see also Figures 2A, C). At E17.5 in the dorsomedial cortex, vast majority of the labeled cells were located in the primitive cortical zone (PCZ), which is the most superficial part of the CP (E, H). In the dorsolateral cortex, most of the labeled cells were still migrating in the CP (E, I). At E18.5 in the dorsomedial cortex, labeled cells were distributed not only in the PCZ, but also in the slightly deeper part of the CP as NeuN-positive mature neurons (F, H). In the dorsolateral cortex, majority of the labeled cells were located in the PCZ (F, I). At P0.5 in the dorsolateral cortex, many labeled cells were distributed in the slightly deeper part of the CP as NeuN-positive mature neurons (G). Small double headed arrows show PCZ. DAPI, 4’,6-diamidino-2-phenylindole; FT, FlashTag; VZ, ventricular zone; MAZ, multipolar cell accumulation zone; IZ, intermediate zone; SP, subplate; CP, cortical plate; PCZ, primitive cortical zone; MZ, marginal zone. Scale bars, 200 μm (A-G), 50 μm (H, I), 10 μm (J).

In summary, labeled cells showed similar migratory profile until they enter the IZ in both dorsolateral and dorsomedial cortices. Cells take longer before they pass the SP and enter the CP in the dorsolateral cortex, compatible with our *in vitro* observations in Figure 2 and similar to the E13.5 cohort. This time lag to enter the CP was not caught up until they settle in their final destination. These observations suggest that regional differences in migration of the E14.5 cohort derive from transient sojourning in the IZ below the SP, at least in part.

### E15.5 cohort

Half a day after injection, at E16.0, many of the labeled cells were in the VZ (Figures 8A, B, Figures S7A, S7H, S7I). We observed some cells outside of the VZ (Figures 8B, Figures S7A, I, arrowheads), which were often positive for Pax6. as in E14.5 cohort. Around the pallial-subpallial boundaries (PSB), small number of cells with single long ascending processes with various orientation were scattered (Figure S7J; similar cells were observed in E17.0 cohort and will be analyzed in detail).

**Figure 8.**
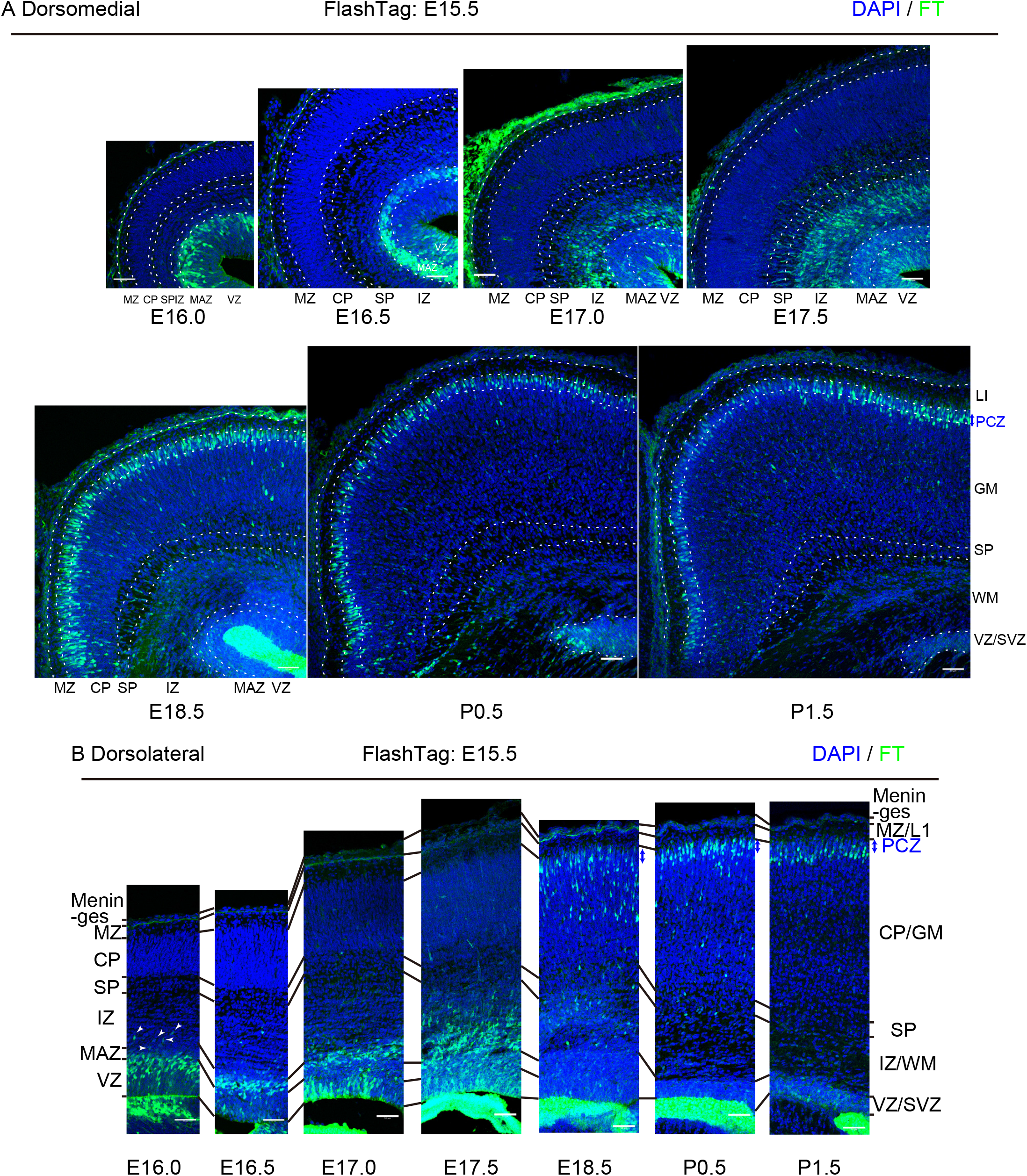
Cohort of cells labeled at E15.5. **A-B:**Coronal sections of E16.0, 16.5, 17.0, 17.5, 18.5, P0.5 and P1.5 brains labeled at E15.5. Data from the dorsomedial cortex and dorsolateral cortex were shown in (A) and (B), respectively. See Figure S7 for low magnification pictures. Figure S7 shows the same sections shown with immunohistochemistry helpful to define histology. At E16.0, most of the labeled cells were in the VZ (A, B, Figures S7A, S7H, S7I). Some labeled cells, often positive for Pax6, were outside of the VZ (B, Figure S7A, S7I) (arrowheads). One day (E16.5; A, B, Figures S7B, S7H, S7I) and 1.5-2 days (E17.0-17.5; A, B, Figures S6C, S6D, S6H, S6I) after injection, most of the labeled cells were in the MAZ and IZ, respectively. At E17.5, most of the labeled cells were migrating in the superficial and deep part of the IZ in the dorsomedial cortex (A, Figures S7D, S7H). In the dorsolateral cortex, migrating cells were mainly in the rather deep part of the IZ (B, Figure S7D, S7I). At E18.5 in the dorsomedial cortex, most of the labeled cells the PCZ (A, Figure S7E, S7H). In the dorsolateral cortex, on the other hand, only the small population of the labeled cells reached the PCZ and others were still migrating in the CP and SP in a locomotion morphology (B, Figures S7E, I). At P0.5, vast majority of the labeled cells settled in the PCZ in the dorsomedial cortex (A, Figures S7F, S7H). In the dorsolateral cortex, too, many labeled cells reached the PCZ (B, Figures S7F, I). At P1.5, labeled cells labeled at E15.5 settled in the gray matter in the dorsolateral cortex (B, Figures S7G, H, I). Some of these labeled cells changed their position slightly apically to leave from the PCZ in the dorsolateral cortex (B, Figures S7G, H, I). DAPI, 4’,6-diamidino-2-phenylindole; FT, FlashTag; VZ, ventricular zone; MAZ, multipolar cell accumulation zone; IZ, intermediate zone; SP, subplate; CP, cortical plate; PCZ, primitive cortical zone; MZ, marginal zone; PSB, pallial-subpallial boundary; LI, cortical layer I (L1 in the IZ is an axonal marker); GM, gray matter; WM, white matter. Scale bars, 50 μm.

One day after injection, at E16.5, most of the labeled cells were accumulated in the MAZ both in the dorsomedial and dorsolateral cortex (Figures 8A, B, Figures S7B, S7H, S7I). 1.5 days after injection, at E17.0, some labeled cells entered the IZ, which is rich in L1-positive axons including thalamocortical and corticofugal axons (Fukuda et al., 1997; Kudo et al., 2005; Yoshinaga et al., 2012), both in the dorsomedial and dorsolateral cortex (Figures 8A, B, Figures S7C, S7H, S7I). Two days after injection, at E17.5, most of the labeled cells were migrating in the superficial and deep part of the IZ in the dorsomedial cortex (Figure 8A, Figures S7D, S7H), but in the dorsolateral cortex, migrating cells were mainly located in the deep part of the IZ (Figure 8B, Figures S7D, S7I).

Three days after injection, at E18.5, most of the labeled cells were located in the PCZ in the dorsomedial cortex (Figure 8A, Figures S7E, S7H). On the other hand, in the dorsolateral cortex, only a small population of the labeled cells reached the PCZ and others were still migrating in the CP and the SP in a locomotion morphology (Figure 8B, Figures S7E, S7I). In the dorsolateral cortex, it took one more day for most of them to reach the PCZ at P0.5 (Figure 8B, Figure S7F, S7I).

At P1.5, cells labeled at E15.5 settled in the gray matter both in the dorsomedial and dorsolateral cortices (Figures 8A, B, Figures S7G, S7H, S7I). In the dorsolateral cortex, some of these labeled cells changed their position slightly deeper to leave the top of the CP, which was not prominently observed in the dorsomedial cortex (Figures 8A, B, Figures S7G, S7H, S7I).

These observations suggest that there are mediolateral differences in migratory profiles in the E15.5 cohort similar to those observed for the E13.5 and E14.5 cohorts.

### E17.0 cohort

Half a day after injection, at E17.5, labeled cells were mainly in the VZ (Figure 9C, Figure S8A). One to 1.5 days after injection, at E18.0-18.5, the main population of the labeled cells migrated out of the VZ into the MAZ (Figures 9C, D, Figures S8B, S8C). They entered the IZ two days after injection, or at E19.0 (Figure 9C, Figure S8D). In the dorsal part of the IZ/WM at P1.0, we observed a band like zone where cellular density is somewhat higher than the deeper and more superficial part of the IZ/WM (Figure 9A, inset). This slightly dense cellular zone in the IZ/WM was sandwiched by L1-positive axon bundles that are skew. At this timepoint, some of the labeled cells were found in this cellular zone in the dorsal part of the IZ/WM (Figure 9A). We also observed small number of labeled cells with single leading processes extending medially (Figure 9A, Figure S8J). Most of the labeled cells at this stage were positive for neuronal marker Hu (Figures S8J2-6). As late as P2.0, or four days after injection, labeled neurons began to migrate in the CP/GM in a bipolar morphology (Figure 9C, Figure S8F). About five or more days after injection, or later than P3.0, labeled cells settle in the PCZ, or the top of the gray matter, of the dorsal cortex (Figure 9C, Figure S8G, H). These cells obtain pyramidal morphology and became positive for NeuN by P5.0 (Figure S8I), suggesting that they were indeed mature neurons.

**Figure 9.**
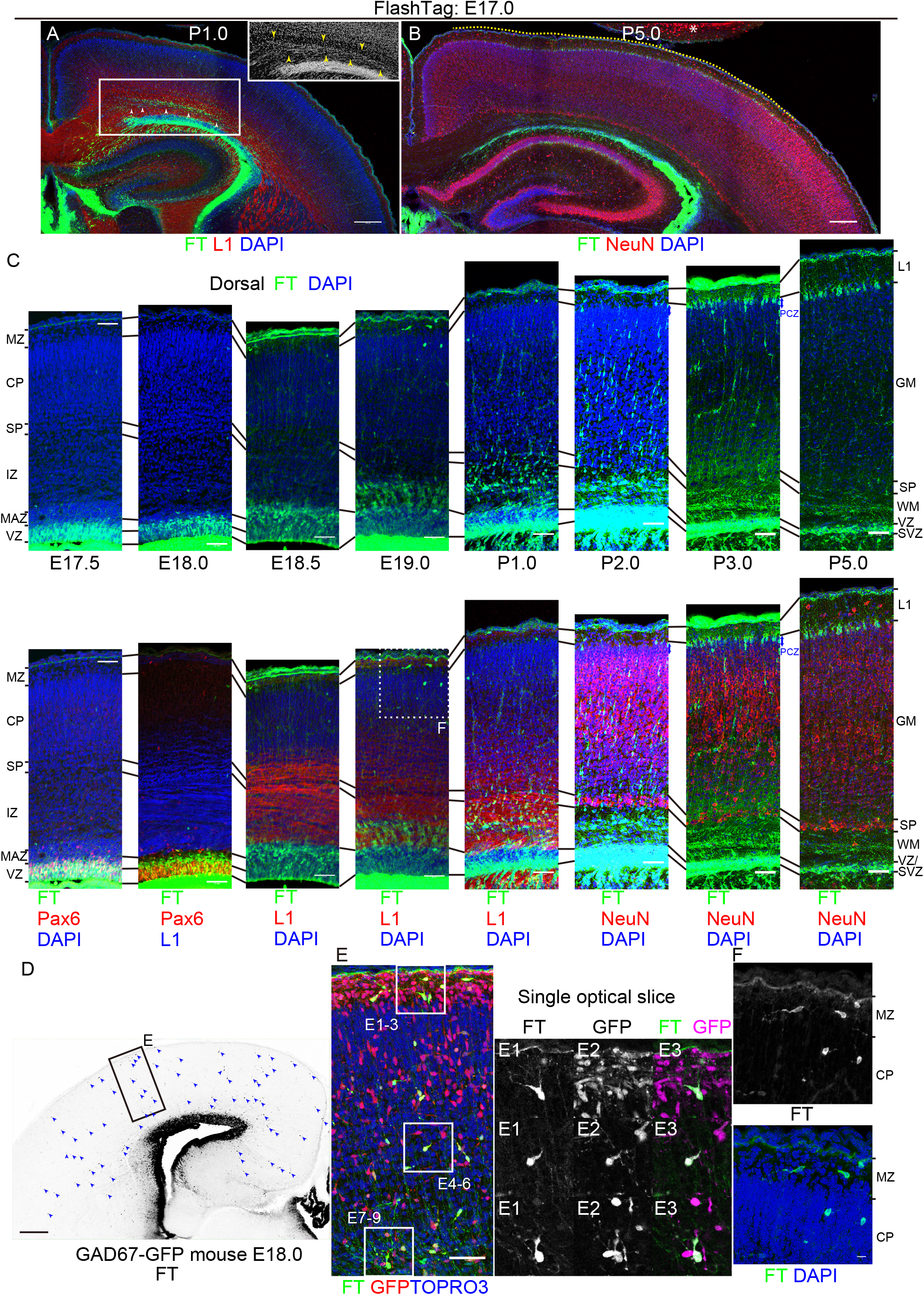
Cohort of cells labeled at E17.0. **A-C:** Coronal section of P1.0 (A) and P5.0 (B) brains labeled at E17.0. See also Figure S8 for lower magnification pictures of E17.5 through P5. Higher magnification pictures of E17.5 through P5 from the dorsal cortex were shown in (C). At E17.5, most of the labeled cells were located in the VZ (C). At E18.0, most of the labeled cells were located in the VZ and MAZ (C). Small number of labeled cells were also found throughout the cortex sparsely. At E18.5, many labeled cells were in the MAZ (C). Some labeled cells sparsely distributed throughout the cortex. At E19.0, many cells entered the L1-positive IZ dorsally (C). Small number of cells were also found in the MZ and CP (F). At P1.0, many labeled cells were migrating in the IZ / white matter (A, C). Migrating cells formed a slightly dense cellular structure (inset in A) sandwiched by L1-positive axon bundles (arrowheads in A). At P2.0, many neurons were migrating in the CP/cortical gray matter with a locomotion morphology (C). At P3.0, many labeled cells reached the dorsal PCZ (C). At P5.0, most of the labeled cells were located in the most superficial part of the cortical gray matter (B, C). Note that many cells were located in the dorsal (and dorsolateral) cortex (the yellow dotted line), and few cells were located in the dorsomedial and lateral cortices. **D-F**: Analyses of GABAergic interneurons. Cells labeled with FT (CytoTell Blue) at E17.0 sparsely distributed throughout the cortex at E18.0 (D, E), and they were mostly positive for GFP in GAD67-GFP mice (E). Labeled cells with similar morphologies were found in the MZ / Layer I and CP at E19.0 before the main population of the labeled cells reach the CP (F). DAPI, 4’,6-diamidino-2-phenylindole; FT, FlashTag; VZ, ventricular zone; MAZ, multipolar cell accumulation zone; IZ, intermediate zone; SP, subplate; CP, cortical plate; PCZ, primitive cortical zone; MZ, marginal zone; PSB, pallial-subpallial boundary; LI, cortical layer I; GM, gray matter; WM, white matter. Scale bars, 200 μm (A, B, D), 50 μm (C, E), 10 μm (F). * indicated another brain on the same slide glass.

Collectively, labeled cells mainly distributed dorsally, and only a few cells settled in the dorsomedial and lateral cortex. We did not observe clear sojourning just below the dorsal and dorsolateral SP as in E13.5-15.5 cohort, but appearance of a slightly dense zone that consists of migrating neurons may suggest sojourning and/or deceleration in the midst of migration in the IZ/WM. Axon bundles just above this zone may contain axons from the SP (Figure 9A, S8E, positive for Nurr1 and Cplx3).

Half a day after injection, at E17.5, some strongly labeled cells with a long ascending process were scattered around the PSB (Figure S8A) as for the E15.5 cohort (Figure 7J). As early as one day after injection, at E18.0, these cells distributed throughout the cortex (Figure 9D, Figure S8B). This population was mostly negative for a radial glial marker Pax6 (Figure 9C, Figures S8A, S8B), glial lineage markers Gfap, Sox10 (Stolt et al., 2002; Zhou et al., 2000) nor Olig2 (Tatsumi et al., 2018) (except for the ventromedial cortex) (data not shown). They were, however, positive for GFP in Gad67-GFP mice (Figure 9E), suggesting that they were GABAergic interneurons. Some of these cells were positive for CGE-derived interneuron markers *Htr3a* (Murthy et al., 2014) and Couptf2 (Kanatani et al., 2015; Kanatani et al., 2008), while others were negative, suggesting that they constitute a heterogenous population. In addition, a few of them were positive for BrdU administered at E13.5, suggesting that at least some of these cells underwent final mitosis days before E17.0. These observations raise a possibility that the FT-labeled interneurons that leave the VZ earlier than the main population of FT-labeled cells are not labeled with FT at mitosis but are labeled after they become postmitotic. Labeled cells with similar morphologies were found in the MZ / Layer I and CP before the main population of the labeled cells reached the CP (Figure 9F, Figures S8C, S8D, S8E).

### Mechanisms of regional differences in neuronal migration

We finally sought to gain insight into the mechanisms of regional differences in neuronal migration. We focused on the sojourning just below the SP in the E14.5 cohort. Based on our observations in Figure 2, we hypothesized that the SP neurons or some other structures in the SP transiently decelerate the migration of later-born neurons in the dorsolateral cortex. First, to see if the SP neurons regulate migration of neurons born at E14.5, we used *reeler* mice, in which SP neurons that normally position below the CP are mispositioned above the CP as revealed by Nurr1 staining (Figures 10A, B) (Hoerder-Suabedissen et al., 2009; Ozair et al., 2018; Pedraza et al., 2014). FT labeling was performed at E14.5 and harvested the brains at E16.5. Compared to wildtype, the mediolateral migratory differences were less clear in *reeler* mice (Figures 10A, B). Next, to see if the thalamocortical axons (Bicknese et al., 1994; Molnár et al., 1998), one of the most characteristic structures in the SP (Figure 10C1), regulate neuronal migration, we used a Crispr/Cas9-based improved-Genome editing via Oviductal Nucleic Acids Delivery (i-GONAD) (Gurumurthy et al., 2019; Ohtsuka et al., 2018; Takabayashi et al., 2018) to generate *Gbx2* knockout mice (Figures S9A-A9C), which lack thalamocortical axons (Hevner et al., 2002; Miyashita-Lin et al., 1999). *Gbx2* knockout mice provide great opportunity to study the role of thalamocortical axons in regulating migration of cortical neurons specifically, because *Gbx2* is expressed in the dorsal thalamus but not in the cortex (Miyashita-Lin et al., 1999). As expected, immunoreactivity against a thalamocortical axon marker Netrin G1 (Nakashiba et al., 2002; Vue et al., 2013) was almost absent in the homozygous mice (Figure 10D1). In these mice, we performed FT labeling at E14.5 and harvested at 50 hours later. FT-labeled migrating neurons showed a migration profile almost identical to that of the control brains (Figures 10C, D), suggesting that thalamocortical axons are not likely to regulate neuronal migration around the SP. Taken together, these observations are compatible with a notion that the SP neurons or some other structures in the SP transiently decelerate the migration of later-born neurons in the dorsolateral cortex, although further analyses are warranted on other structures in the subplate and cell-autonomous regulation of neuronal migration that is dependent on cortical regions.

**Figure 10.**
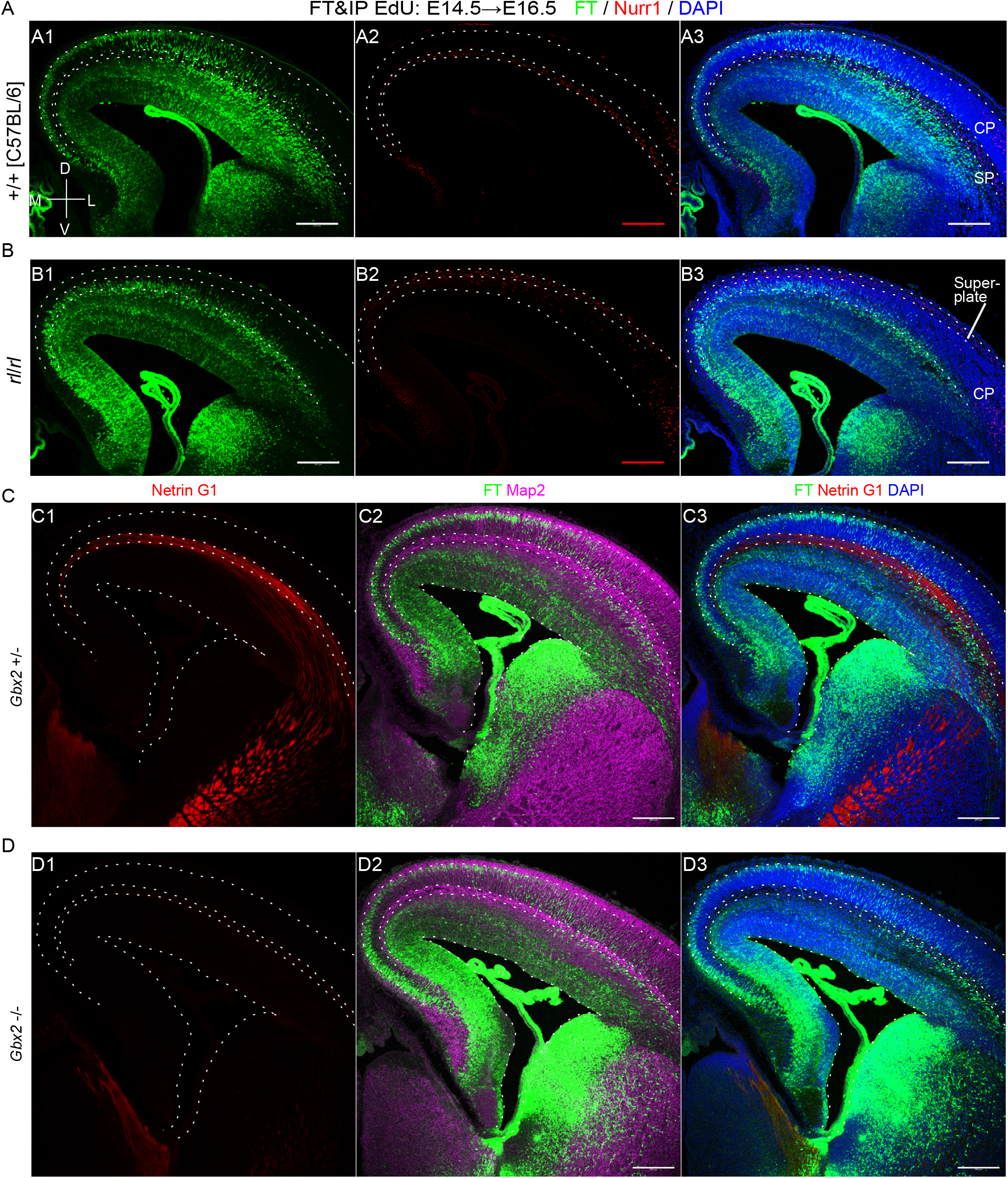
Regional differences in neuronal migration in *reeler* mutants and *Gbx2* -/- mice. **A-B:** In wildtype brains, Nurr1+ cells were observed in the SP (A), while in *reeler* mice, Nurr1+ cells were mostly observed in the superplate, or beneath the meninges (B). In contrast to wildtype mice clearly showing regional differences in neuronal migration (A), regional differences were not clear in *reeler* mice (B). FT was performed at E14.5 and fixed at E16.5. **C-D:** In *Gbx2* +/- brain, Netrin G1-positive thalamocortical axons run the SP (C1). In *Gbx2* -/- brain, Netrin G1-positive thalamocortical axons were almost absent in the cortex (C2). In both cases, many neurons were observed just beneath the SP. FT was performed at E14.5 and fixed 50 hour later. Coronal sections slightly caudal to the main part of the interventricular foramina were shown to evaluate the thalamus at the same sections. DAPI, 4’,6-diamidino-2-phenylindole; FT, FlashTag; VZ, ventricular zone; MAZ, multipolar cell accumulation zone; IZ, intermediate zone; SP, subplate; CP, cortical plate; M, medial; L, lateral; D, dorsal; V, ventral. Scale bars, 200 μm.

## Discussions

Using FT technology, we showed that there are clear regional differences in neuronal migration in the pallium even where there is an underlying VZ. The regional differences were dependent on the embryonic stages when the apical radial glial cells divide at the ventricular surface to produce neuronal progenitors and neurons. In E10.5 and E11.5 cohorts, regional differences in neuronal migration, which is defined in the current study as movement from mitosis at the ventricular surface to settlement just beneath the meningeal surface, were not clear. In E12.5 cohort, we described slight regional differences. In E13.5, E14.5 and E15.5 cohorts, neurons in the dorsomedial cortex reached the top of the CP about one day earlier than those in the dorsolateral cortex. In E17.0 cohort, we observed that labeled neurons positioned nearly only dorsally. We also observed migratory behavior of the subpopulation of the labeled cells, for example, mitotically-active, Pax6 positive cells that leave the VZ as early as 0.5 days after labeling in E14.5 cohort (Figures 7A, I, J, Figure S6A). These comprehensive descriptions provide basic information about cortical development.

How are the regional differences formed? Timelapse imaging suggested that cells labeled at E14.5 in the dorsolateral cortex stop transiently below the SP while those in the dorsomedial cortex do not. It is known that the SP neurons interact with later-born neurons (Ohtaka-Maruyama et al., 2018). In *reeler* cortex, in which SP neurons are superficially mispositioned below the meninges and migrating neurons do not make contact with SP before entering the CP, mediolateral migratory difference was not clear. This observation was compatible with the idea that the SP cells transiently decelerate the migration of later-born neurons as part of normal migration. In addition to the SP neurons and thalamocortical axons, there are many other structures potentially relevant to the migratory difference; corticofugal (Denaxa et al., 2001), catecholaminergic (Lidov and Molliver, 1982) axons, and radial fibers bending (Mission et al., 1991; Saito et al., 2019) and branching (Takahashi et al., 1990) around the SP. We cannot, in addition, exclude the possibility that cell-intrinsic mechanisms of migratory neurons are also involved. Further research, including *in vivo* transplantation and specific ablation of anatomical structures, would be needed to obtain the mechanistic insight.

What is the physiological role of the regional differences in migratory profiles? This regional difference of migratory profiles of the E14.5 cohort was clearly visualized with FT but less clearly with thymidine analogues, a standard approach to study neurogenic gradient. We thus think that this has a biological significance different from neurogenic gradient. Migrating neurons receive synaptic contacts from the SP neurons when they pass the SP (Ohtaka-Maruyama et al., 2018). At the same stage, thalamocortical fibers wait in the SP (Lopez-Bendito and Molnar, 2003). Thus, if migrating neurons slow down beneath the dorsolateral SP, they have more chance to interact with the SP and/or thalamic afferents. Along the developmental time axis, the regional difference in neuronal migration including sojourning beneath the SP, was clear in cohorts that contain future layer IV neurons. Histologically, the dorsomedial cortex, where labeled cells did not sojourn beneath the SP clearly, is agranular and lacks layer IV. The dorsolateral cortex, where cells sojourn just beneath the SP, corresponds to primary somatosensory areas, where layer IV neurons are predominant. These observations suggest a possibility that sojourning beneath the SP might be implicated in thalamocortical circuit formation and/or layer IV formation. In line with this, role of the extracellular environment is estimated to be increasingly important in refining neuronal identity as they migrate and differentiate especially in the E14-labeled future layer IV neurons (Telley et al., 2019). In addition, abnormal migration and positioning of neurons labeled at E14.0 (mainly future layer IV neurons) results in abnormal differentiation (Oishi et al., 2016a; Oishi et al., 2016b).

PP splitting involves the establishment of the cortical plate within the PP (Goffinet and Lyon, 1979; Marin-Padilla, 1971). It has been assumed that the cells in the earliest CP are future layer VI cells and that their active reorganization drives PP splitting (Nichols and Olson, 2010; Olson, 2014). However, it is also possible that some future SP neurons actively migrate away. In our present study of earliest cohorts, labeled cells were first observed in the PP, then in the CP and MZ upon the formation of the CP, and finally they moved down below the CP, supporting the downward movement of some SP neurons through the CP. This observation is compatible with previous descriptions using timelapse imaging or *in vivo* observations in which future SP neurons are labeled with *in utero* electroporation (Saito et al., 2019), genetically (*Lrp12*/*Mig13a*-*EGFP* mice) (Schneider et al., 2011) and immunohistochemically (Hpca+/Reelin- and Eaac1+/Reelin-)(Osheroff and Hatten, 2009). Historical studies using ^3^H-TdR in cats (Luskin and Shatz, 1985) described that future SP neurons are transiently located in the deep part of the histologically-defined CP (Boulder-Committee, 1970) although Luskin and Shatz assumed that this is part of the SP. Altman and Bayer (Bayer and Altman, 1991) analyzed rats using ^3^H-TdR and suggested that the SP neurons temporarily reside in the CP. These observations suggest that at least some neurons in the earliest CP and MZ are future SP neurons. Since FT might label only subpopulation of cells in our study, however, we do not exclude a possibility that some earliest born neurons form a distinct cell layer below the CP before or right after the CP formation.

Neurons labeled at E17.0 mainly distributed dorsally and relatively rarely medially nor laterally (Figure 9B). However, small number of cells labeled at E17.0 were observed at E18-19 in dorsomedial cortex as well. Some of them might migrate to subiculum and hippocampus; others fan out sparsely in the cingulate and secondary motor cortices. Another possibility is that they divide abventricularly to lose fluorescence. Around this stage, gliogenesis accelerates, but it was also shown that there are many Hopx-positive neurogenic moRG in the medial cortex at this stage (Vaid et al., 2018). Still another possibility is that they undergo programmed cell death. The fate of the majority of the dorsomedial E17.0 cohort remains to be determined.

Interneurons are born in the ganglionic eminences and preoptic area. Ventral progenitors were also labeled with FT, but FT-labeled neurons rarely entered the cortex. This can be explained by the frequent abventricular division in the ventral forebrain (Katayama et al., 2013; Tan et al., 2016; Tan and Shi, 2013) (Figures S1D, S1E). However, small number of cortical GABAergic interneurons were labeled on E17 (Figures 9D, E) and presumably on E15.5 (Figure S7J), which distributed into the cortex within a day. One interpretation for this retaining of the label in interneurons in E15.5 and E17 cohorts is that FT potentially labels a certain subpopulation that undergo final mitosis relatively late for interneurons at ventricular surface. Another interpretation is labeling of migrating interneurons that undergo final mitosis earlier than dye injection. Ventricle-directed migration of interneurons, some of which touch the ventricular surface, was described from E13 in mice and E15 in rats (Nadarajah et al., 2002). If some of the migrating interneurons had touched the ventricular surface and been labeled with FT, however, FT-labeled interneurons should have been observed in our earlier cohort as well. In our E14.5 cohort, we indeed observed some cells with long ascending processes that left the VZ earlier than the main population (Figures 7I, J). However, they were mostly positive for Pax6, a dorsal progenitor marker, suggesting that they are a different population. The origin of these interneurons labeled with FT on E15.5 and E17 remains to be determined.

In the E14.5-15.5 cohorts, we observed cells that left the VZ within 0.5 days mainly in the dorsolateral cortex. These cells have the distribution, migratory behavior, cycling feature and morphology in common with “rapidly exiting population,” or REP, which was labeled by *in utero* electroporation and subsequent BrdU incorporation (Tabata et al., 2009). Many of these cells were positive for radial glial markers Pax6 and Sox2 (Figure 7J), supporting the view that moRG cells comprise a subpopulation of REP (Tabata et al., 2012). In the earlier cohort (labeled at E12.5, Figure 5G, Figures S4G, S4H), we also observed similar Pax6-positive population that left the VZ early. This population, however, did not share the lateral-more to medial-less gradient of the distribution of REP that we previously reported. In early stages of corticogenesis when the CP is not formed yet, migrating neurons show multipolar morphology (Hatanaka et al., 2004; Tabata and Nakajima, 2003) but they do not accumulate just above the VZ to form a clear MAZ. Because the accumulated multipolar cells were shown to serve as a fence to limit the apical border of the range of interkinetic nuclear migration (Watanabe et al., 2018), these Pax6-positive cells in our E12.5 cohort that leave the VZ soon and distribute sparsely may result from a poor fence to limit the apical border of the VZ. Conversely, we suppose that the REP and moRG in later cohorts (E14.5 and 15.5) may have an active mechanism to pass the MAZ/SVZ.

Application of FT to visualize neuronal migration has several strengths over conventional methods. First, FT has potential to detect differences in neuronal migration that cannot be detected by thymidine analogues. Second, this method enables visualization of neuronal migration of the whole brains. This feature especially goes well with whole brain 3D approach including FAST (Seiriki et al., 2017; Seiriki et al., 2019). Third, the methodology is simple, and FT can be a versatile approach to study neuronal migration in the whole brain in healthy and disease model mice. On the other hand, FT has several technical limitations. First, tangential migration of projection neurons (e.g. lateral dispersion of the rostromedial telencephalic wall-derived future SP neurons (Pedraza et al., 2014), ventral streaming of pallial-derived, early embryonic preplate neurons (Saito et al., 2019) and abnormal tangential migration of projection neurons (Pinheiro et al., 2011)) could not be efficiently visualized because FT labels mitotic cells on the ventricular surface throughout the brain. Second, the migration profile might be biased toward the slowly exiting population (SEP) (Tabata et al., 2009), or direct progeny of apical progenitors, because the fluorescence of the secondary proliferative population would decrease upon mitosis.

The results of FT experiments must be interpreted considering the following limitations. First, one may think that the mediolateral migratory difference is not true regional difference but a simple reflection of medially thinner cortex. But this is less likely because the regional difference was preserved in the posterior cortex, where the thickness of the cortical wall is equivalent in the dorsomedial and dorsolateral cortex (Figure 2E, Supplemental movie. 1). Second, we measured developmental stages using embryonic days, but developmental stages are confounded by neurogenic gradient, which differs about one day mediolaterally (Takahashi et al., 1999). This view is especially important to discuss the regional difference in neuronal migration of the E13.5 cohort. Cells labeled at E13.5 reached just beneath the meningeal surface at about E14.5-15.0 in the dorsomedial cortex upon the formation of the CP (Figures 6B, C, F). At this stage, somal translocation was previously observed (Nadarajah et al., 2001). It is also reported that multipolar cells do not transform into bipolar locomotion cells before the CP forms (Hatanaka et al., 2004). In the dorsolateral cortex at this stage, in contrast, the CP structure is already formed, and labeled neurons need to transform from a multipolar migration mode to a bipolar locomotion mode (Figures 6C, D, G). One may reason that this difference in migratory modes determine the mediolateral differences in neuronal migration profiles. However, this is unlikely because the mediolateral regional difference was also observed in E14.5 and 15.5 cohort, when the dorsomedial CP is well developed.

In summary, we applied FT to describe neuronal migration and described migratory profiles of early- and late-born projection neurons in normal mouse cortical development. The labeling features of FT shed light into the hitherto overlooked regional differences of neuronal migration profiles. This versatile approach would be useful to study neuronal migration of disease models and transgenic animals.

## Acknowledgements

This work was supported by Grants-in-Aid for Scientific Research of the Ministry of Education, Culture, Sports, Science, and Technology (MEXT)/Japan Society for the Promotion of Science Grants-in-Aid for Scientific Research (KAKENHI) (JP17J05365, JP18K19379, JP19H05227, JP18K07855, JP19H01152, JP19K08306, JP20H03649, JP16H06482, JP18K19378, JP20H00492, JP19H05217, JP18H05416), the Keio Gijuku Academic Development Funds, Keio Gijuku Fukuzawa Memorial Fund for the Advancement of Education and Research, Takeda Science Foundation, and PRIME, AMED (JP19gm6310004, JP20dm0207061). S.Y. was a Research Fellow of Japan Society for the Promotion of Science from fiscal year (FY) 2017 to FY 2019.

We thank Drs. Ludovic Telley and Denis Jabaudon (University of Geneva) for technical advice and valuable discussions. We also thank Core Instrumentation Facility, Collaborative Research Resources, Keio University School of Medicine, Dr. Yoshifumi Takatsume, and distinguished technicians including Emiko Shimeno, Miki Sakota, Noriko Suzuki, Chisa Konno, and Maiko Saito for technical assistance. Greatest gratitude is expressed to all the members of Nakajima Lab for the valuable advice, expertise and encouragement.

## Author Contributions

Conceptualization, S.Y., K.K. and K.N.; Methodology, S.Y. M.T., A. Kasai., and H.H.; Investigation, S.Y., M.K., A. Kitazawa, K.I., and M.T.; Writing ––– Original Draft, S.Y.; Writing ––– Review & Editing, K.K. and K.N.; Visulalization, S.Y., M.T., and A. Kasai; Supervision, K.K. and K.N.; Funding Acquisition, S.Y., A. Kasai, H.H., K.K., and K.N.

## Declaration of Interests

The authors declare no competing interests.

## Supplemental Figure Legends

**Figure S1**, related to Figure 1

Characterization of cell population labeled with FT.

A: The definition of “dorsomedial”, “dorsal” and “dorsolateral” cortex in the current study. B: The schematic presentation of histological zones in this study. See the *histological terminology* section in the materials and methods for discussion. C: CytoTell Blue was injected into the LV of the E12.5 GAD67-GFP brains. A coronal section of E15.5 brains slightly caudal to the section shown in Figure 1M was shown. In the “reservoir” (Altman & Bayer, 1991), there were many migrating cells that were mostly negative for GFP (C1-C3). More ventrally, labeled cells were identified in the caudal amygdaloid stream (CAS), and were negative for GFP (C4-C6). Arrowheads show rare examples of cells positive for both FT and GFP. D-E. Immunohistochemistry against pH3 was performed in E13.5 (D) and E14.5 (E) wild type brains in which CFSE was injected at E12.5. Abventricular mitosis labeled by pH3 was abundant in the ganglionic eminences (D, E). In the medial ganglionic eminence (MGE) at E13.5, many FT-labeled cells were observed in the VZ and apical half of the SVZ (D). At E14.5, when FT-labeled interneurons enter the cortex when fluorescent dyes were injected into the parenchyma of the ganglionic eminences (data not shown), FT-labeled cells were again observed in the VZ and apical half of the SVZ (E). Note that relatively small number of cells migrated in the deep part of the SVZ of the MGE and in the presumptive pallidum, and that few labeled cells with interneuron-like morphology was observed in the cortex. F. CytoTell Blue was injected to parenchyma of the ganglionic eminence (GE) of the heterozygous GAD67-GFP mice at E12.5. Asterisk in (F) indicates the injection site retrospectively identified. Strongly labeled cells distributed in the whole hemispheres, especially in the SVZ and marginal zone (MZ) (F, F1). They often showed tangential morphology and were positive for GFP (F1-F10).

DAPI, 4’,6-diamidino-2-phenylindole; FT, FlashTag; GAD-GFP, Glutamate decarboxylase 67-green fluorescent protein, VZ, ventricular zone; MAZ, multipolar cell accumulation zone; IZ, intermediate zone; SP, subplate; CP, cortical plate; PCZ, primitive cortical zone; MZ, marginal zone; LI, cortical layer I; GM, gray matter; WM, white matter; PSB, pallial-subpallial boundary; MGE, medial ganglionic eminence; LGE, lateral ganglionic eminence; Ctx, cortex. Scale bars, 200 μm (C, D, E, F), 50 μm (F1), 20 μm (C1-6).

**Figure S2**, related to Figure 3

Cohort of cells born at E10.5.

A-F: Coronal sections of 11.5 (A, A’), 12.5 (B), 13.5 (C), 14.5 (D, D’), 15.5 (E, E’) and 16.5 (F) brains labeled at E10.5. See also Figure 3 for higher magnifications. As early as E11.5, some cells were found in the preplate (PP), which was very thin in the dorsomedial cortex, as well as in the VZ (A, A’). At E12.5, many cells were in the PP (B). In the dorsomedial cortex at E13.5, labeled cells were in the PP (C). In the dorsolateral cortex, on the other hand, many labeled cells were located in the CP and MZ (C). At E14.5, thin CP was identified in the dorsomedial cortex as well (D, D’). Some labeled cells were in the deep part of the CP in the dorsomedial cortex, but many labeled cells were in the MZ (D, D’). In the dorsolateral cortex, many labeled cells were found near the boundary between the CP and SP (D, D’). At E15.5, labeled cells were found at the boundary between SP and CP as well as MZ in the dorsomedial cortex (E), which is similar to the dorsolateral cortex of the E14.5 (D). In the E15.5 dorsolateral cortex, many labeled cells were in the CSPG-positive SP (E, E’). At E16.5, in both the dorsomedial and dorsolateral cortex, labeled cells were mainly found in the SP (F). Some cells were also found in the MZ (F). G: At E16.5, in both the dorsomedial and dorsolateral cortex, labeled cells were mainly found in the SP and were Tbr1-positive. Some cells were also found in the MZ and were positive for Reelin, suggesting that they were Cajal-Retzius neurons.

DAPI, 4’,6-diamidino-2-phenylindole; FT, FlashTag; SP, subplate; CP, cortical plate; MZ, marginal zone. Scale bars, 200 μm (A-F) and 50 μm (G).

**Figure S3**, related to Figure 4

Cohort of cells born at E11.5.

A-F: Coronal sections of E12.0 (A), E12.5 (B), E13.0 (C), E13.5 (D), E14.5 (E) and E15.5 (F) brains labeled at E11.5. See the legend for Figure 4 for explanation. G-H: Immunohistochemistry against Ctip2 and CSPG of E15.5 brains in which FT was performed at E11.5. Images (G) (dorsomedial) and (H) (dorsolateral) were taken from insets in Figures 4D and E, respectively. At E15.5, many cells were in the lower part of CP and, to lesser extent, MZ. Some cells were also found in the SP in the dorsolateral cortex. Most of the labeled cells in the CP at E15.5 were positive for Ctip2, a deep layer marker.

DAPI, 4’,6-diamidino-2-phenylindole; FT, FlashTag; SP, subplate; CP, cortical plate; MZ, marginal zone. Scale bars, 200 μm (A-F), 20 μm (G, H).

**Figure S4**, related to Figure 5

Cohort of cells born at E12.5.

**A-F:** Coronal sections of E13.0 (A), 13.5 (B), 14.0 (C), 14.5 (D), 15.5 (E), and 16.5 (F) brains labeled at E12.5 shown with nuclear staining. See the legend for Figure 5 for explanation. **G-H:** Single optical slices of E 13.0 brains taken from the dorsomedial and dorsolateral cortices were shown in (G) and (H), respectively. In the dorsomedial cortex at E13.0, many labeled cells were in the VZ, but a small number of labeled cells were also found in the PP (Figure 5A, G). The latter cells were often weakly positive for Pax6 (G, arrowheads). In the dorsolateral cortex, many labeled cells were located in regions just above the VZ in addition to the VZ, and they are often negative for Pax6 (H; arrows). FT+ / Pax6+ cells outside of the VZ were relatively rare (H, an arrowhead).

DAPI, 4’,6-diamidino-2-phenylindole; FT, FlashTag; PP, preplate; VZ, ventricular zone; MAZ, multipolar cell accumulation zone. Scale bars, 200 μm (A-F) and 10 μm (G, H).

**Figure S5**, related to Figure 6

Cohort of cells born at E13.5.

**A-E:** Coronal sections of E14.0 (A), 14.5 (B), 15.0 (C), 15.5 (D) and 16.5 (E) brains labeled at E13.5 and stained with DAPI. See the legend for Figure 6 for explanation. **F-G:** Coronal sections of E17.5 (A) and 18.5 (B) brains labeled at E13.5. At E17.5, many strongly labeled neurons located in the rather deep part of the CP in the dorsomedial and dorsolateral cortices (Figure 6). Even at this stage, in the most lateral part of the cortex, many labeled cells were still migrating radially or about to leave the reservoir (R) (Bayer and Altman, 1991) (F1). Labeled cells were also found in the caudal amygdaloid stream (CAS) (F2). At E18.5, labeled cells were distributed in the CP but not in the SP (G).

DAPI, 4’,6-diamidino-2-phenylindole; FT, FlashTag; Pir, piriform cortex; Ins, insular cortex. Scale bars, 200 μm (A-F, F1, G), 50 μm (F2).

**Figure S6**, related to Figure 7

Cohort of cells born at E14.5.

**A-G:** Coronal sections of E15.0 (A), 15.5 (B), 16.0 (C), 16.5 (D), 17.5 (E), 18.5 (F) and P0.5 (G) brains labeled at E14.5. Images for FT and nuclear staining are shown. See also the legend of Figure 7 for explanation. **H:** A coronal section of a P7 brain in which FT labeling and intraperitoneal BrdU injection was performed at E14.5. In the dorsolateral cortex, FT labeled cells mainly distributed in the layer IV. In dorsomedial and lateral cortex, FT labeled cells mainly distributed in the layer II/III. BrdU positive cells were mainly detected in the superficial layers.

DAPI, 4’,6-diamidino-2-phenylindole; FT, FlashTag; VZ, ventricular zone; MAZ, multipolar cell accumulation zone; IZ, intermediate zone; SP, subplate; CP, cortical plate; PCZ, primitive cortical zone; MZ, marginal zone. Scale bars, 200 μm.

**Figure S7**, related to Figure 8

Cohort of cells labeled at E15.5.

**A-J:** Coronal sections of E16.0 (A), 16.5 (B), 17.0 (C), 17.5 (D), 18.5 (E), P0.5 (F) and P1.5 (G) brains labeled at E15.5. Higher magnification pictures from the dorsomedial cortex and dorsolateral cortex were shown in (H) and (I), respectively. Higher magnification of the pallial-subpallial boundaries (PSB) of E16.0 (0.5 day after injection) brains was shown in (J). See Figure 8 for legends. At E16.0, around the pallial-subpallial boundary (PSB), small number of labeled cells were in the IZ and CP with a long leading process (J).

DAPI, 4’,6-diamidino-2-phenylindole; FT, FlashTag; VZ, ventricular zone; MAZ, multipolar cell accumulation zone; IZ, intermediate zone; SP, subplate; CP, cortical plate; PCZ, primitive cortical zone; MZ, marginal zone; PSB, pallial-subpallial boundary; LI, cortical layer I (L1 in the IZ is an axonal marker); GM, gray matter; WM, white matter. Scale bars, 200 μm (A-G), 50 μm (H-J).

**Figure S8**, related to Figure 9

Cohort of cells labeled at E17.0.

**A-I:** Coronal section of E17.5 (A), 18.0 (B), 18.5 (C), 19.0 (D), P1.0 (E), P2.0 (F), P3.0 (G) and P5.0 (H) brains labeled at E17.0. Higher magnification pictures from these brains are shown in Figure 9C. At E17.5, most of the labeled cells were located in the VZ (A). Some of the labeled cells scattered in the brain parenchyma (arrowheads in A). At E18.0, most of the labeled cells were located in the VZ and MAZ (B). Again, small number of labeled cells distributed throughout the cortex (arrowheads in B). At E18.5, many labeled cells were in the MAZ (C). Some labeled cells sparsely distributed throughout the cortex. At E19.0, many cells entered the L1-positive IZ dorsally (D). At P1.0, many labeled cells were migrating in the IZ (E). Migrating cells were migrating in a zone sandwiched by L1-positive axon bundles (Figure 9A). This zone was deeper than the SP, as visualized by Nurr1 and Cplx3, SP neuron markers. At P2.0, many neurons were migrating in the CP/cortical gray matter with a bipolar morphology (F). At P3.0, many labeled cells reached the dorsal PCZ (G). At P5.0, most of the labeled cells were located in the most superficial part of the cortical gray matter (H), and were positive for NeuN (I). **J:** At P1.0, labeled cells were migrating dorsally, as well as ventrally (to the hippocampus) and, to lesser extent, medially. Some labeled cells were also found in the VZ/SVZ. Most of these were positive for a neuronal marker Hu (J2-J6).

DAPI, 4’,6-diamidino-2-phenylindole; FT, FlashTag; IZ, intermediate zone; SP, subplate; CP, cortical plate. Scale bars, 200 μm (A-H, J1), 50 μm (J2-3), 20 μm (J4-6), 10 μm (I).

**Figure S9**, related to Figure 10

Generation of *Gbx2* knockout mice using Crispr/Cas9

**A:** A schematic diagram of our strategy to make a deletion of a region which contains a homeobox domain. Key concepts of this strategy were based on mice generated in a previous study (Wassarman et al., 1997). Animals with an allele in which a region between Target 1 and Target 2 was deleted were screened by a band shift in electrophoresis of the PCR products, and further confirmed by sequencing. **B-C:** Direct sequencing of the PCR products amplified from the G0 mice (B, Target 1; C, Target 2).

**Supplemental movie 1**, related to Figure 2E

Whole-brain imaging of migrating neurons. FT labeling was performed at E14.5, fixed about two days later and stained the nuclei with Hoechst. The movie shows series of coronal sections i) in the posterioanterior (PA) direction from the occipital pole of the cortex to the frontal pole (FT only), ii) back in the AP direction slightly past the level of interventricular foramen (FT + Hoechst), iii) in the PA direction to the presumptive frontal cortex (FT only), and iv) in the AP direction slightly past the interventricular foramen (FT + Hoechst).

## Materials and Methods

### KEY RESOURCES TABLE

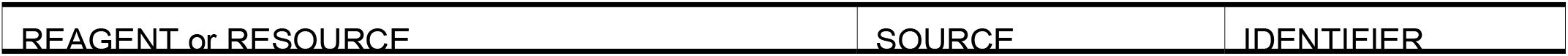
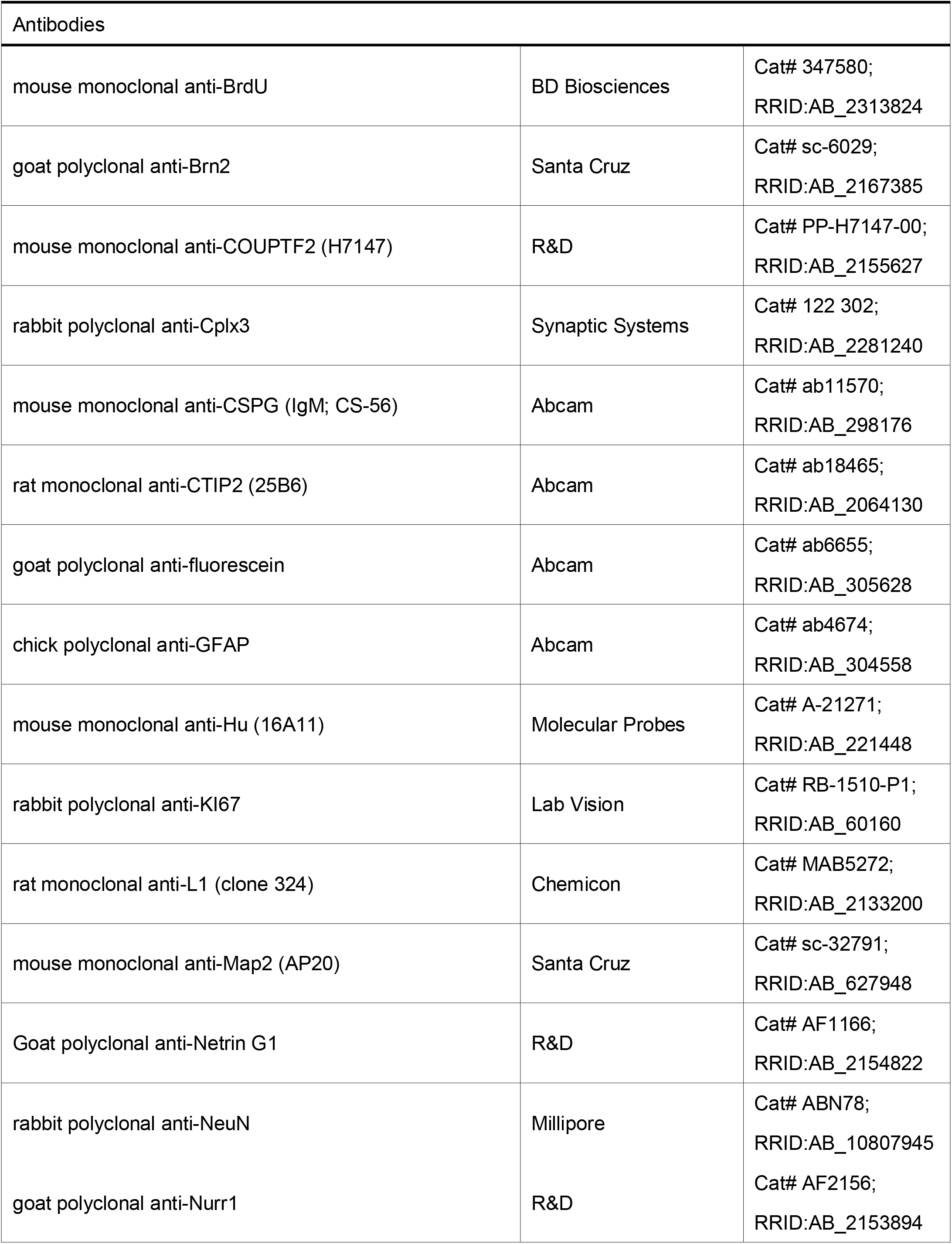
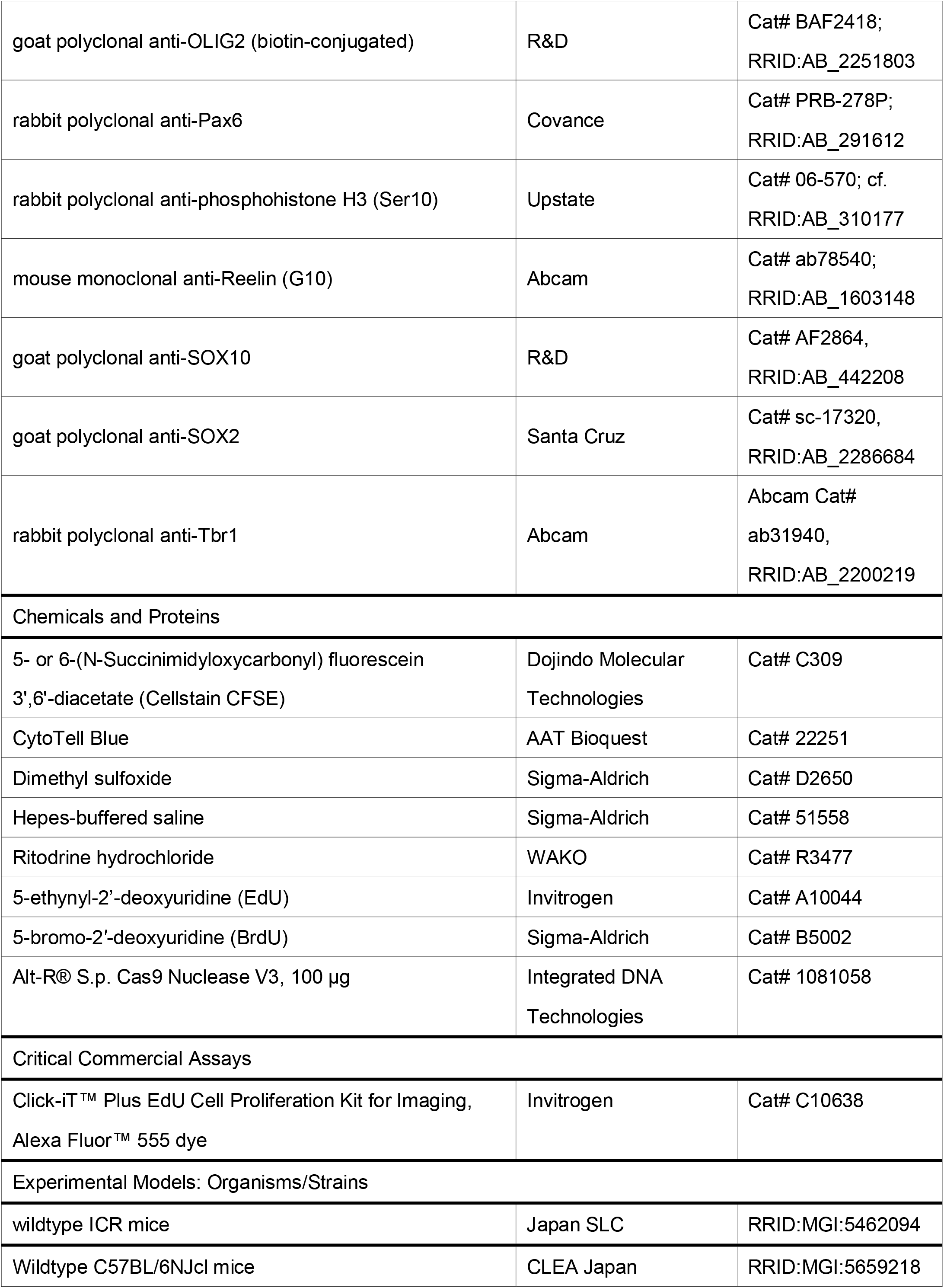
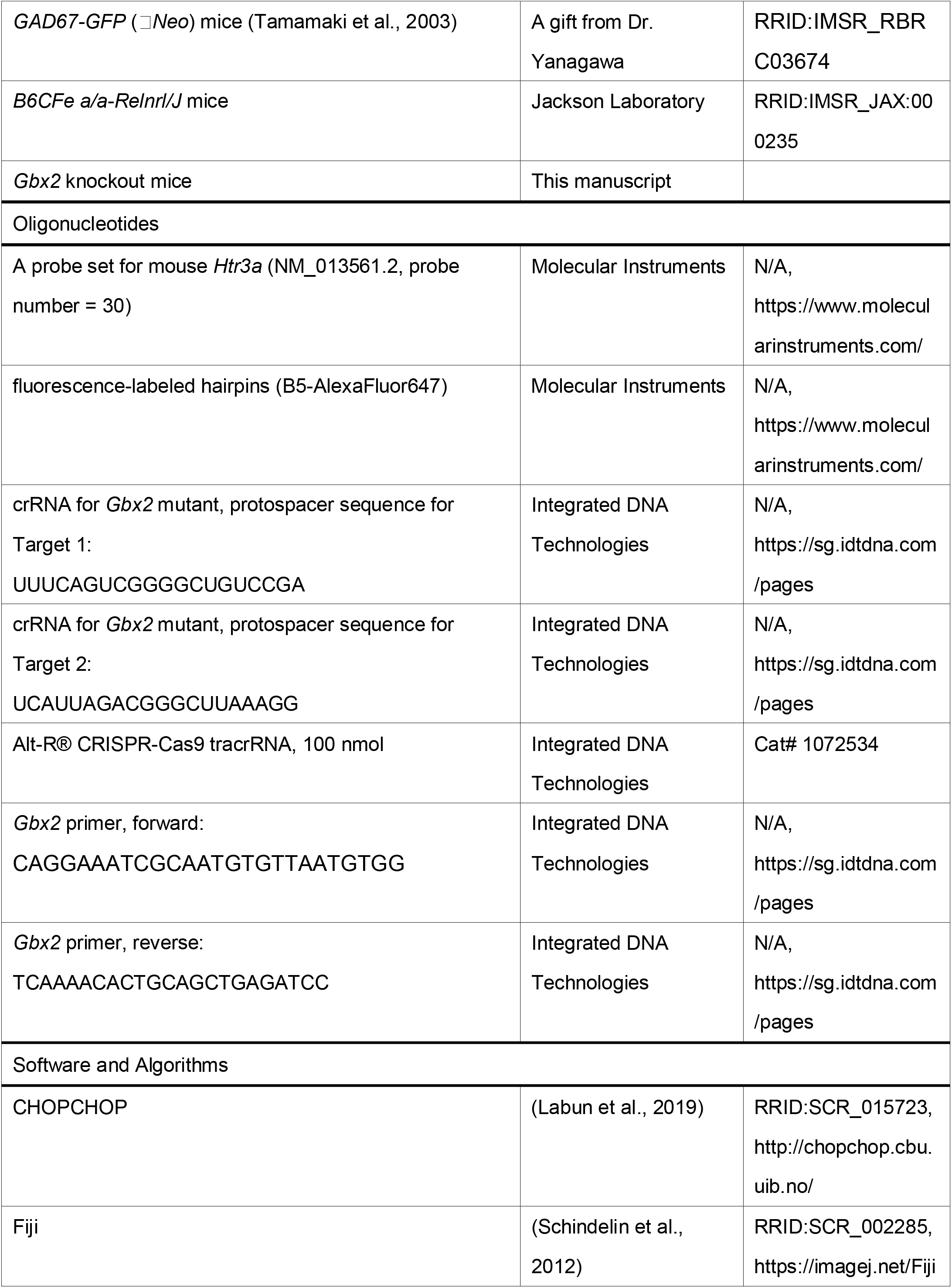
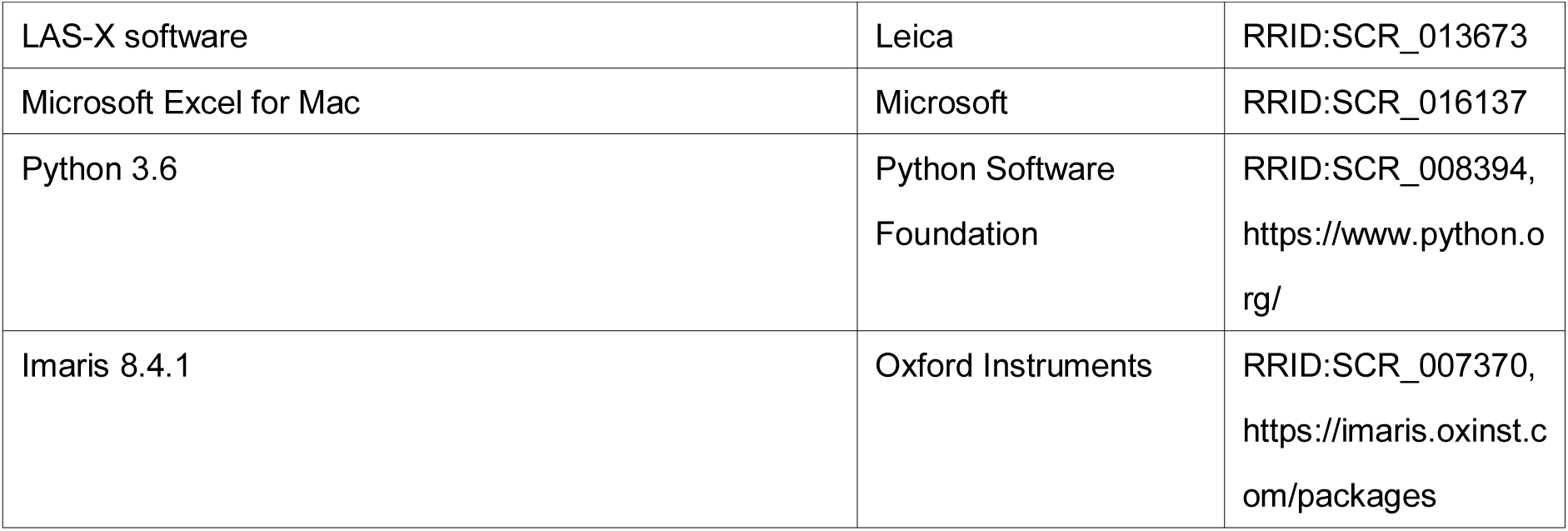

#### Lead Contact

Further information and requests for resources and reagents should be directed to and will be fulfilled by the Lead Contact, Kazunori Nakajima.

### RESOURCE AVAILABILITY

#### Materials Availability

This study did not generate new unique reagents. We did not deposit the *Gbx2* knockout mice generated in the present study, because they were designed based on a mouse line *Gbx2*^tm1Mrt^ (MGI:3665450) used in a previous study (Wassarman et al., 1997). Requests for the mice should be directed to the Lead Contact.

#### Data and Code Availability

This study did not generate/analyze datasets or code.

## EXPERIMENTAL MODEL AND SUBJECT DETAILS

### Animals

Pregnant wildtype ICR (RRID:MGI:5462094) and C57BL/6NJcl mice (RRID:MGI:5659218) were purchased from Japan SLC (Shizuoka, Japan) and CLEA Japan (Tokyo, Japan). *GAD67*-*GFP* (ΔNeo) mice (Tamamaki et al., 2003) were provided by Dr. Yanagawa (Gunma University, Gunma, Japan), and heterozygous progenies were backcrossed to wild type ICR mice. Heterozygous males were mated with wild type ICR mice and used in the experiments. *Reeler* mice (B6CFe a/a-*Reln*^*rl*^/J; RRID:IMSR_JAX:000235) were obtained from the The Jackson Laboratory and maintained by mating heterozygous females with homozygous males. The day on which a vaginal plug was detected was considered embryonic day (E) 0. Dams, pups, and weaned animals were kept under a 12/12-hour light/dark cycle in a temperature-controlled room. The animals had free access to food and water. Embryos and pups of both sexes were indiscriminately analyzed, because sexes cannot be macroscopically determined. All animal experiments were performed according to the Institutional Guidelines on Animal Experimentation at Keio University.

## METHOD DETAILS

### FT Surgical procedures

Pregnant mice were deeply anesthetized, and their intrauterine embryos were manipulated as described previously (Nakajima et al., 1997). FT was performed as described as previously (Telley et al., 2016) with some modification. 10 mM 5- or 6-(N-Succinimidyloxycarbonyl) fluorescein 3’,6’-diacetate (Cellstain CFSE, C309, Dojindo Molecular Technologies, Inc., Kumamoto, Japan) working stock was prepared by dissolving CFSE in Dimethyl sulfoxide (DMSO) (Hybri-Max™, Sigma-Aldrich, St. Luis, MO). The working solution was further diluted with 1X Hepes-buffered saline (HBS) to make 1 mM solution just before surgery. The solution was colored with Fast Green (final concentration 0.01-0.05%) to monitor successful injection. In experiments using GAD67-GFP mice, CytoTell Blue (22251, AAT Bioquest, Sunnyvale, CA) was used instead of CFSE. About 0.5 μl of the prepared FT solution was injected into the lateral ventricle. Trans-illumination method (Shimogori and Ogawa, 2008) was utilized to visualize small embryos of E10.5 and 11.5. At these early stages, 200 μl of 0.1 mg/ml Ritodrine hydrochloride (WAKO, now FUJIFILM Wako Pure Chemical Corporation, Osaka, Japan) was injected intraperitoneally to relax myometrium (Nishiyama et al., 2012; Takeo, 2016; Takeo et al., 2015). After applying plenty of phosphate-buffered saline (PBS) into the abdominal cavity and onto the surface of manipulated uterine horns, injected embryos were placed back into the abdominal cavity.

### Administration of thymidine analogues

EdU and BrdU (Sigma) was dissolved in PBS at 5 mg/mL and 10 mg/mL, respectively. Bolus intraperitoneal injection of EdU or BrdU solution was performed at 25 μg/g body weight (BW) and 50 μg/gBW, respectively.

### Histological terminology

The VZ and SVZ was determined according to the definition provided by Boulder’s Committee (Boulder-Committee, 1970). Because the VZ is a pseudostratified columnar epithelium, their nuclei, by definition, are mostly oriented radially. The basal border of the VZ nuclei was also able to be determined by a radial glial marker Pax6 staining (Englund et al., 2005) or acute administration of a thymidine analogue (Tabata et al., 2012) because the nuclei of radial glia in the S-phase occupies a basal zone of the VZ (interkinetic nuclear migration). Just above the VZ is a zone that we previously named the MAZ (Tabata et al., 2009), where multipolar cells that have just exited the VZ transiently accumulate. The cell density of this zone is high, and nuclei are randomly oriented (Bayer and Altman, 1991; Yoshinaga et al., 2012). Although many cells in the MAZ are postmitotic (Tabata et al., 2009), there are some cycling cells in the MAZ. The MAZ and the lower part of the SVZ, which is originally characterized by abventricular cells with proliferative activity by Boulder’s Committee, overlaps. Just above the MAZ is a zone rich in L1-positive axonal fibers (Yoshinaga et al., 2012) and somata of the immature migrating neurons. We called this zone the IZ according to the Boulder’s Committee’s suggestion. The SP layer, which was described after Boulder’s Committee defined histological terminology, was excluded from the IZ in the current study, because the main component is relatively mature subplate neurons. The original description by the Boulder Committee defined the IZ and SVZ as distinct regions, but because we observe many proliferative cells in the axon-rich area (Tabata et al., 2009; Vaid et al., 2018) (Figures 1A-D), the IZ in our definition and the SVZ inevitably overlaps. Collectively, the SVZ starts from the MAZ extending into the IZ in our definition. We therefore preferred the use of the MAZ and IZ to describe neuronal migration more precisely, except for contexts stressing abventricular mitosis. We defined the SP according to the cytoarchitechtonic criteria and presence of abundant CSPG (Bicknese et al., 1994). The CP was determined by cytoarchitectonic criteria (high cellularity, radial orientation of the nuclei (Olson, 2014)) and/or weak immunoreactivity of CSPG (Bicknese et al., 1994). The primitive cortical zone, or PCZ (Sekine et al., 2011; Shin et al., 2019), was determined by weak or lack of NeuN staining in the CP. The marginal zone was determined by cytoarchitectonic criteria––most superficial, hypocellular zones just above the CP. Before the formation of the CP, the zone between proliferative zone (i.e. the VZ at this stage) and the meningeal surface was named as the PP (Bystron et al., 2008), although this area might include intermediate progenitors (Vasistha et al., 2015) as well. Cytoarchitecture changes as development proceeds, which is summarized in Figure S1B.

The definition of dorsomedial, dorsal and dorsolateral cortex in the coronal sections was provided in Figure S1A. We obtained images at the rostrocaudal axis of foramina of Monro unless otherwise specified. We obtained dorsolateral high magnification images from regions the lateral borders of which cross the pallial-subpallial angles. We obtained dorsomedial high magnification images from regions adjacent to the medial protrusion of the lateral ventricles. In most of cases the images were corrected so that the apicobasal axes are parallel to a line that passes medial protrusion of the lateral ventricles and ipsilateral pallial-subpallial angels. In the late stages of cortical development, the dorsomedial high magnification images were shown with dorsal-up because lines that pass medial protrusion of the lateral ventricles and ipsilateral pallial-subpallial angels are no longer parallel to the apicobasal axes nor perpendicular to the meningeal surfaces.

### Histological sample preparation

The harvested embryonic brains were fixed by immersing in 4% paraformaldehyde (PFA) at 4°C with gentle agitation for 1 hour to overnight. The postnatal embryos were perfused with ice-cold 4% PFA, and their brains were further fixed by immersing in 4% paraformaldehyde (PFA) at 4°C with gentle agitation for several hours to overnight. The brains were cryoprotected by immersing in 20% and 30% sucrose in PBS at 4°C for several hours to overnight sequentially, embedded in 75% O.C.T. compound (Sakura, Tokyo, Japan) (O.C.T: 30% sucrose = 3:1) and frozen with liquid nitrogen. Brains were cryosectioned coronally by 20 μm thick on MAS-coated slide glass (MAS-02; Matsunami Glass Ind.,Ltd., Osaka, Japan)

For immunohistochemistry, sections were immersed with PBS with 0.01% Triton X-100 (Sigma-Aldrich, St. Louis, MO) (PBS-Tx) for more than 30 minutes at room temperature (RT). Antigen retrieval was performed in most of the experiments by incubating in 1x HistoVT ONE (NACALAI TESQUE, INC., Kyoto, Japan) at 70°C for 20 minutes. To detect BrdU, sections were treated with a sodium citrate buffer (pH 6) at 105°C for 5 minutes and with 2 M hydrogen chloride at 37°C for 30 minutes. The sections were blocked with 10% normal goat serum in PBS-Tx at RT, and incubated with the primary antibody overnight at 4°C. After washing with PBS-Tx for three times, the sections were incubated with secondary antibodies for 1 hour at RT. The details of the primary antibodies are shown in Key Resource Table and the antibody characterization section.

Histological detection of EdU was performed using Click-iT™ EdU Cell Proliferation Kit for Imaging, Alexa Fluor™ 555 dye (C10338, Thermo Fisher Scientific) according to the manufacturer’s protocol.

When nuclear staining was performed without immunohistochemistry, sections were immersed with PBS for more than 30 minutes at RT and incubated with 2.5 ng/μl of 4’,6-diamidino-2-phenylindole (DAPI; D3571; Thermo Fisher Scientific, Waltham, MA) or 0.5 μM of TO-PRO3 Iodide (T3605, Thermo Fisher Scientific) at RT for 1 hour. When nuclear staining was performed with immunohistochemistry, DAPI was added to the secondary antibody solution. Sections were mounted using PermaFluor (TA-030-FM; Thermo Fisher Scientific).

### Antibody Characterization

The antibodies used in this study were listed in Key Resource Table. The mouse anti-BrdU antibody (clone B44) (Tabata et al., 2009) was used to detect nuclei of cells that were in the S phase when BrdU was administered. This antibody is derived from hybridization of mouse Sp2/0-Ag14 myeloma cells with spleen cells from BALB/c mice immunized with iodouridine-conjugated ovalbumin (manufacturer’s datasheet and a previous report (Gratzner, 1982)). This antibody detects BrdU (but not thymidine) in single-stranded DNA, free BrdU, or BrdU coupled to a protein carrier. The antibody also reacts with iodouridine, which was not used in this study.

An anti-Brn2 antibody was used as a layer II/III/V marker (Oishi et al., 2016a). This antibody was raised against a peptide mapping at the C-terminus of *BRN2* of human origin (manufacturer’s datasheet). Although previous study suggested that this antibody detects both Brn1 and Brn2 in western blotting (Yamanaka et al., 2010), we believe that this antibody predominantly detects Brn2 in immunohistochemistry of perinatal cortical slices, because electroporation of a shRNA against *Brn2* significantly diminished immunoreactivity of this antibody but not of anti-Brn1 antibody while electroporation of a shRNA against *Brn1* did not significantly diminish immunoreactivity of this antibody in immunohistochemistry (Oishi et al., 2016a). Even if this antibody detects Brn1 as well, its expression pattern is similar to that of Brn2 in the developing cerebral cortex and the use of this antibody as a layer marker would be justified.

An anti-COUP-TF II antibody was used to label CGE- and PoA-derived interneurons (Kanatani et al., 2015; Kanatani et al., 2008). This mouse monoclonal antibody was raised against recombinant human COUP-TF II (amino acids 43-64) (manufacturer’s datasheet). The specificity of this antibody was previously confirmed by absence of immunohistochemical staining in a Couptf2 conditional knockout tissue (Suh et al., 2006). An anti-Cplx3 antibody was used to label the SP in the postnatal stage (Hoerder-Suabedissen et al., 2009). This antibody was raised against recombinant mouse Complexin3 (amino acids 1-158) (manufacturer’s datasheet). The specificity of this antibody was previously confirmed by absence of signals in a *Cplx3* knockout tissue in immunohistochemistry and western blotting (Reim et al., 2009).

An anti-CSPG antibody was used to label the PP, MZ and SP (Bicknese et al., 1994). This antibody was well characterized elsewhere (Yi et al., 2012).

Ctip2/Bcl11b was used as a deep layer marker (Arlotta et al., 2005). Anti-CTIP2 rat monoclonal antibody was raised against a fusion protein corresponding to human CTIP2 (amino acids 1-150). This antibody detects 2 bands representing Ctip2 at about 120kD (manufacturer’s datasheet). This antibody detected nuclear staining in wildtype mice while no signals in *Ctip2*-null mice on immunohistochemistry (Zhang et al., 2012).

A goat anti-fluorescein antibody was used to boost FT signals when brains were analyzed days after FT injection and fluorescent labeling was weak. This antibody was raised against fluorescein conjugated to goat IgG. Western blotting detected BSA conjugated fluorescein (manufacturer’s datasheet). This antibody enhanced fluorescence from FT-labeled cells, but no signal was detected in untreated (CFSE was not injected) brains (data not shown).

A mouse anti-Hu monoclonal antibody was used as a neuronal marker (Marusich et al., 1994; Tabata and Nakajima, 2003). This antibody was raised against a human HuD peptide (QAQRFRLDNLLN-C)-Keyhole Limpet Hemocyanin conjugate, and recognizes HuC, HuD and HuDpro in western blotting (Marusich et al., 1994). This antibody showed immunoreactivity similar to human anti-Hu autoantibody in western blotting of human neuron extract, which was blocked by synthetic HuD peptide (Marusich et al., 1994).

A rabbit anti-Ki-67polyclonal antibody was used to label proliferating cells. This antibody was raised against a synthetic peptide from the human Ki-67 protein. Immunohistochemistry of human lymph nodes resulted in nuclear staining of germinal center (manufacturer’s datasheet). Proliferating reactive astrocytes (Chen et al., 2017) and colorectal carcinoma foci (Zhao et al., 2017) were reported to be specifically labeled. Immunohistochemistry of developing mouse cortex resulted in nuclear staining of the proliferative zones including VZ and SVZ, as previously published (Watanabe et al., 2018).

L1 immunohistochemistry was performed to label the IZ rich in axons including thalamocortical and corticofugal axons (Fukuda et al., 1997; Kudo et al., 2005; Yoshinaga et al., 2012). The antibody used was raised against glycoprotein fraction from cerebellum of 8-10 day old C57BL/6J mice. The same clone from the previous vendor did not stain fiber bundles in the *L1*-null mice (Fransen et al., 1998).

An anti-Map2 monoclonal antibody [AP20] was used as a subplate marker (Ohtaka-Maruyama et al., 2013; Ohtaka-Maruyama et al., 2018). This antibody was raised against cow MAP-2 (amino acids 997-1332), and detects bands corresponding to MAP2A/B on western blotting (manufacturer’s datasheet). Immunohistochemistry of developing mouse cortex resulted in an identical staining pattern previously reported with other antibody against MAP2 (AB5622; Merck Millipore) (Ohtaka-Maruyama et al., 2013; Ohtaka-Maruyama et al., 2018).

Netrin G1 immunohistochemistry was used to mark thalamocortical axons (Nakashiba et al., 2002). The anti-Netrin G1a antibody was raised against purified insect cell line *Sf* 21-derived recombinant mouse Netrin-G1a (rmNetrin-G1a) (manufacturer’s datasheet). Mouse Netrin-G1a specific IgG was purified by mouse Netrin-G1a affinity chromatography. Manufacturer’s datasheet states that this antibody shows less than 2% cross-reactivity with rmNetrin-1, rchNetrin-2 and rhNetrin-4. Cortical immunoreactivity was lost in *Gbx2* conditional knockout mice, in which thalamocortical axons failed to innervate (Vue et al., 2013).

NeuN immunohistochemistry was used to label neuronal cells. The anti-NeuN antibody used was an affinity purified rabbit polyclonal antibody raised against GST-tagged recombinant mouse NeuN N-terminal fragment (ABN78, Millipore) (manufacturer’s datasheet). This antibody is a rabbit polyclonal version of anti NeuN antibody (mouse monoclonal, MAB377, Millipore, clone A60), and has been widely used as a neuronal marker by authors of many different literatures [e.g. (Ataka et al., 2013; Huang et al., 2015; Lundgaard et al., 2015).] Immunohistochemistry of *Rbfox3/NeuN*-null tissue using this antibody and the mouse monoclonal antibody (clone A60), which also has been widely used as a neuronal marker and was extensively characterized by western blotting and 2D electrophoresis (Lind et al., 2005), detected no signals (Lin et al., 2018). Double immunohistochemistry using ABN78 and A60 resulted in an identical staining pattern.

A goat anti-Nurr1 antibody was used to label the SP neurons. This antibody was raised against *E. coli*-derived recombinant mouse Nurr1 (Val332-Lys558) (manufacturer’s datasheet), and reported to detect the nuclei of the SP neurons (Hoerder-Suabedissen et al., 2009; Ozair et al., 2018; Pedraza et al., 2014). No signal was detected in *Nurr1*-deficient mice (data not shown).

A rabbit polyclonal anti-phospho-histone H3 antibody (Ser10) (06-570, Upstate, Spartanburg, SC) was used to label mitotic cells (Hendzel et al., 1997; Kim et al., 2017). This antibody was raised against a short peptide from the amino-terminus of H3 from amino acids 7-20 (A7RKSTGGKAPRKQL20C) synthesized containing a single phosphorylated serine at position 10 (Hendzel et al., 1997). This antibody detected a single band in whole cell protein and acid-soluble nuclear protein from Colcemid-treated mitotic Hela cells but did not detect in whole cell protein and acid-soluble nuclear protein from interphase enriched preparation (Hendzel et al., 1997).

A rabbit anti-Pax6 antibody was used to label radial glial cells. This antibody was raised against a peptide (QVPGSEPDMSQYWPRLQ) derived from the C-terminus of the mouse Pax-6. Western blotting of mouse Raw264.7 cells detects a single band (manufacturer’s datasheet). In the cerebellum of chimera mice made from wildtype and Pax6-null cells, nuclear immunoreactivity was detected in wildtype granular cells while no signal was detected in Pax6 null cells (Swanson and Goldowitz, 2011). In our study, nuclear immunoreactivity was detected in the majority of the VZ cells (Englund et al., 2005) and small number of extra-VZ cells, as expected (Shitamukai et al., 2011; Vaid et al., 2018).

Reelin was used as a marker for Cajal-Retzius cells (Ogawa et al., 1995). Anti-Reelin monoclonal antibody[G10] was raised against a recombinant fusion protein, corresponding to amino acids 164-496 of Mouse Reelin. This antibody detects an expected 388kDa band on western blotting (manufacturer’s technical information). On immunohistochemistry, this antibody detected Cajal-Retzius cells in the marginal zone in the developing wildtype cortex but no signal was detected in the *reeler* cortex except for blood vessels (Ishii et al., 2019), confirming its specificity.

A goat anti-Sox2 polyclonal antibody was used to label nuclei of radial glia. This antibody is an affinity purified antibody raised against a peptide mapping near the C-terminus of human SOX2. Western blotting of human and mouse embryonic stem cells detected a single band at 34 kDa (manufacturer’s datasheet). Immunohistochemistry of developing mouse (Vaid et al., 2018; Watanabe et al., 2018) and human cortex (Nowakowski et al., 2016) resulted in labeling of radial glial cells in the VZ and SVZ.

Tbr1 has been widely used as a marker for postmitotic neurons of the PP, SP and deep layer (Hevner et al., 2001). The detailed information about the antibody used in the current study was described elsewhere (Betancourt et al., 2014).

A chicken anti-GFAP antibody was used to label astrocytes. This chicken polyclonal IgY antibody was raised against a recombinant full-length protein corresponding to Human GFAP, isotype 1. Western blotting of mouse and rat cortical lysates detected a single band (manufacturer’s datasheet). A number of studies have used this antibody to label astrocytes (Saliu et al., 2014). In GFAP-Cre driven GFP transgenic mice, immunoreactivity from this antibody showed excellent colocalization with GFP signals (Suarez-Mier and Buckwalter, 2015), confirming its specificity.

A goat anti-SOX10 antibody and goat anti-OLIG2 antibody was used to label oligodendrocyte (Stolt et al., 2002; Zhou et al., 2000) progenitors and oligodendrocyte + astrocyte progenitors(Tatsumi et al., 2018), respectively. The anti-SOX10 antibody was raised against *E. coli*-derived recombinant human SOX10 (Met1-Ala118) (manufacturer’s datasheet) and has been widely used to label cells of the oligodendrocyte lineage in many literatures including mouse spinal cord (Kelenis et al., 2018) and dorsal cortex (Winkler et al., 2018) in immunohistochemistry. The anti-OLIG2 antibody used in this study was raised against *E. coli*-derived recombinant human SOX10 (Met1-Ala118). In Western blots, less than 5% cross-reactivity with recombinant human (rh) OLIG1 and rhOLIG3 is observed, according to the manufacturer’s technical information. This antibody has been used to label cells of the glial progenitors on immunohistochemistry (Tabata et al., 2009).

### in situ HCR

Fluorescent *in situ* hybridization was performed using in situ HCR v3.0 (Choi et al., 2018). E18.0 brains in which FT was performed at E17.0 were perfused with ice-cold 4% PFA and post-fixed overnight at 4°C. Brains were embedded in 3% low-melting agarose gel and vibratomed by 100 μm thick. Brain slices were preserved at -20°C in a cryoprotectant solution (30% w/v sucrose, 1% w/v polyvinyl-pyrrolidone (PVP)-40, 30% v/v Ethylene glycol in PBS) until use. Brain slices were washed in PBS for 5 minutes at RT, and incubated in a hybridization solution (Molecular Instruments, Los Angeles, CA) at 37°C in a 96-well plate with agitation (a round shaker, 200rpm). The probe set for mouse *Htr3a* (NM_013561.2, probe number = 30) was designed by and purchased from Molecular Instruments. Brain slices were incubated with 4 nM probes overnight at 37°C with agitation. After washing with a prewarmed wash solution (Molecular Instruments) for 15 minutes three times at 37°C with agitation and with 5x SSC with 0.1% Tween20 (5x SSCT) for 5 minutes three times at RT with agitation, sections were incubated with fluorescence-labeled hairpins (B5-AlexaFluor647) reconstituted with an amplification solution (Molecular Instruments) overnight at RT with agitation. After washing with 5x SSCT for more than 5 minutes three times at RT with agitation and counterstaining with DAPI, brain slices were mounted using PermaFluor on MAS-coated slide glasses. This resulted in essentially the same staining pattern as previously described (Murthy et al., 2014).

### Image Acquisition of glass slide samples

The fluorescence images were acquired through confocal laser scanning microscopes (FV1000; Olympus, Tokyo, Japan & TCS SP8; Leica, Wetzlar, Germany). Stitching was performed with LAS-X software (Leica) (RRID:SCR_013673) equipped with the Leica confocal microscope, when necessary. Images were analyzed with Fiji (RRID:SCR_002285) (Schindelin et al., 2012). Linear changes in tone and background subtraction were made. Maximum projection images of optical slices were made to show the entire morphology of the whole cortical wall. Single optical slices were shown to evaluate colocalization of signals of different channels.

### Whole-brain imaging and generation of 3D image movies

Three-dimensional imaging of a whole brain was performed using block-face serial microscopy tomography (FAST) (Seiriki et al., 2017; Seiriki et al., 2019) with some modification. Briefly, brains were perfused with ice-cold PBS and ice-cold 4% PFA. The harvested brains were post-fixed for one week at 4°C. The fixed brains were stained with Hoechst33258 (Seiriki et al., 2019) and embedded in the previously reported 4% oxidized agarose (Ragan et al., 2012). Subsequently, whole-brain images were obtained at a spatial resolution of 1.0 × 1.0 × 5.0 μm^3^. The resulting section images were stitched by FASTitcher, written in Python 3.6 (Seiriki et al., 2019). We generated 3D-rendered movies from 2D stacks of serial stitched images using Imaris 8.4.1 (Bitplane, Belfast, UK).

### Time-Lapse Analyses

Time-lapse observations of slice culture were performed using a previously described (Tabata and Nakajima, 2003). Briefly, coronal brain slices (200 µm thick) were cultured in Neurobasal medium (NB) containing 2% B27 (Invitrogen) on MilliCell-CM culture plate inserts (PICM03050; Merck KGaA, Darmstadt, Germany). The dishes were then mounted in a CO2 incubator chamber (40% O2, 65% N2, 5% CO2 at 37°C) fitted onto a confocal microscope TCS SP8. Approximately 10-20 optical Z sections were obtained automatically every 30 minutes. Photobleaching was linearly corrected afterward to maintain signal strength of the labeled cells using Fiji to enable visual evaluation of migration profiles.

### i-GONAD

CRISPR guide RNAs were designed using CHOPCHOP (Labun et al., 2019). The synthetic crRNAs, tracrRNA, and Cas9 protein were commercially obtained as Alt-R™ CRISPR guide RNAs from Integrated DNA Technologies (Coralville, IA) and Alt-R™S.p. Cas9 Nuclease V3. Adult ICR mice were purchased from Japan SLC. Females in estrus were mated with stud males. Females used were not superovulated. Surgical procedures for i-GONAD were performed as described previously with minor modifications (Gurumurthy et al., 2019; Ohtsuka et al., 2018) at around E0.7 under deep anesthesia. Mixture of 15 μM gRNA for Target 1, 15 μM gRNA for Target 2, and 1 μg/μl Cas9 protein was prepared in Opti-MEM. 0.02% Fast Green was used to monitor successful injection. Approximately 1.5 μl of electroporation solution was injected into the oviduct from upstream of the ampulla using a glass micropipette. The electroporation was performed using NEPA21 (NEPA GENE, Tokyo, Japan) (poring pulse: 50 V, 5 ms pulse, 50 ms pulse interval, 4 pulses, 10% decay, single pulse orientation and transfer pulse: 10 V, 50 ms pulse, 50 ms pulse interval, 3 pulses, 40% decay, ± pulse orientation). Animals carrying the expected deletion were mated with wildtype ICR mice.

## QUANTIFICATION AND STATISTICAL ANALYSIS

Histological samples were evaluated by visual inspection. Careful anatomical, qualitative analyses were performed in most of the experiments. All quantitative data presented are expressed as arithmetic mean ± SEM, and the exact values of n (number of brains) are provided in the Results section and Figure Legends. Descriptive statistics values, including mean, SEM and percentage, were calculated using Microsoft Excel for Mac (RRID:SCR_016137). No hypothesis tests were conducted.

## Notes

### Competing Interest Statement

The authors have declared no competing interest.

