## Supplementary figures and images for "Comprehensive characterization of migration profiles of murine cerebral cortical neurons during development using FlashTag labeling"

### Figure S1

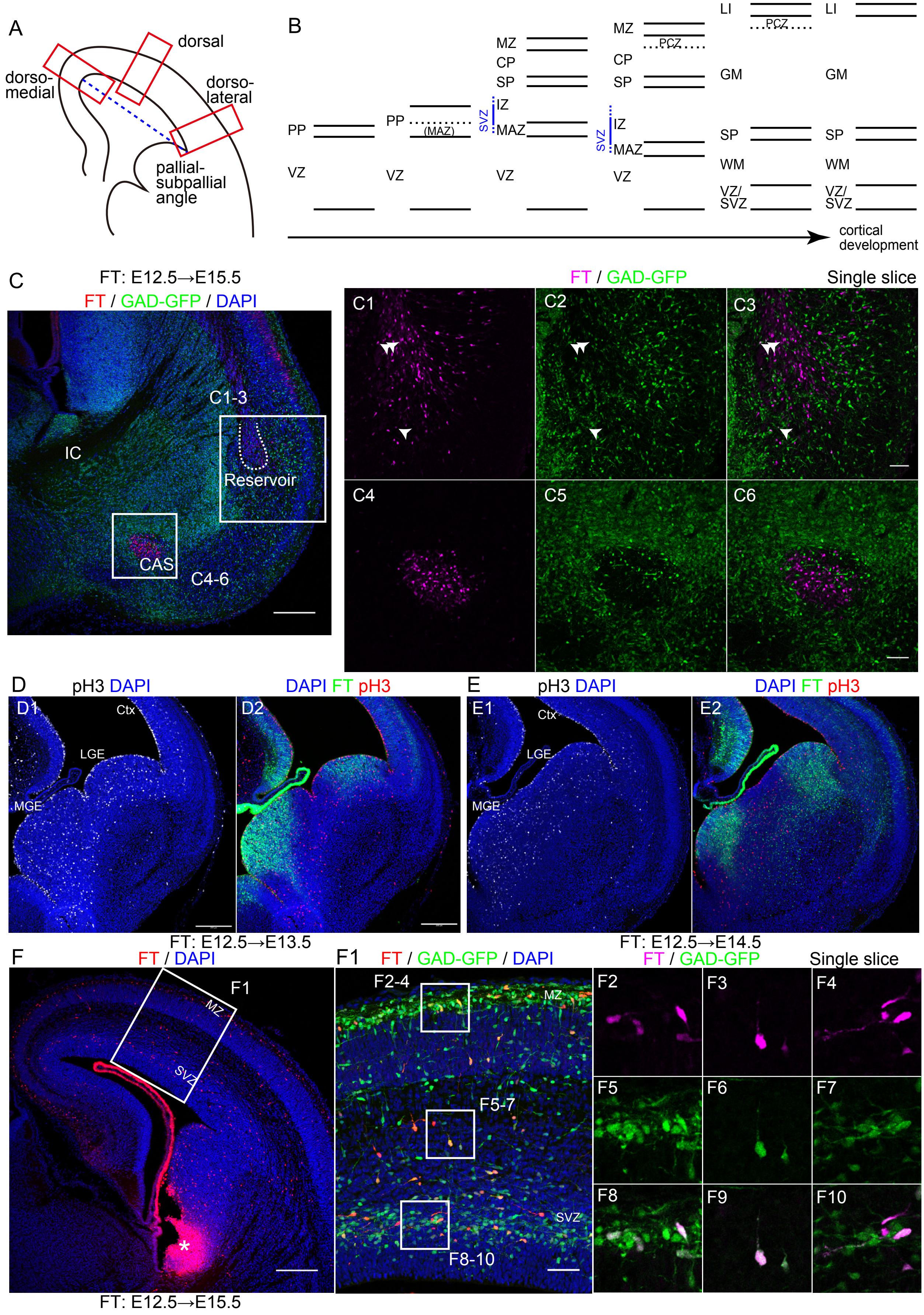

### Figure S2

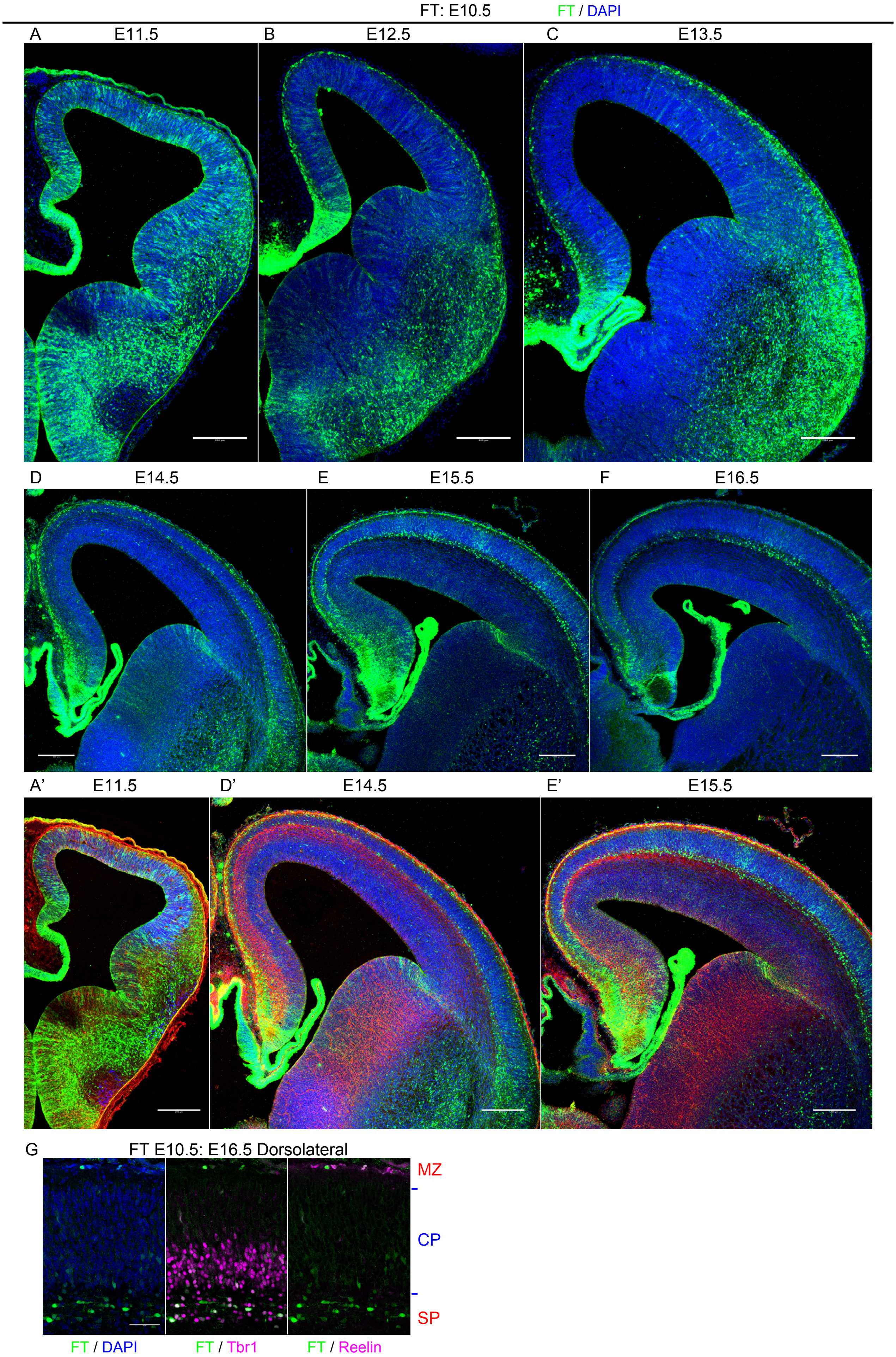

### Figure S3

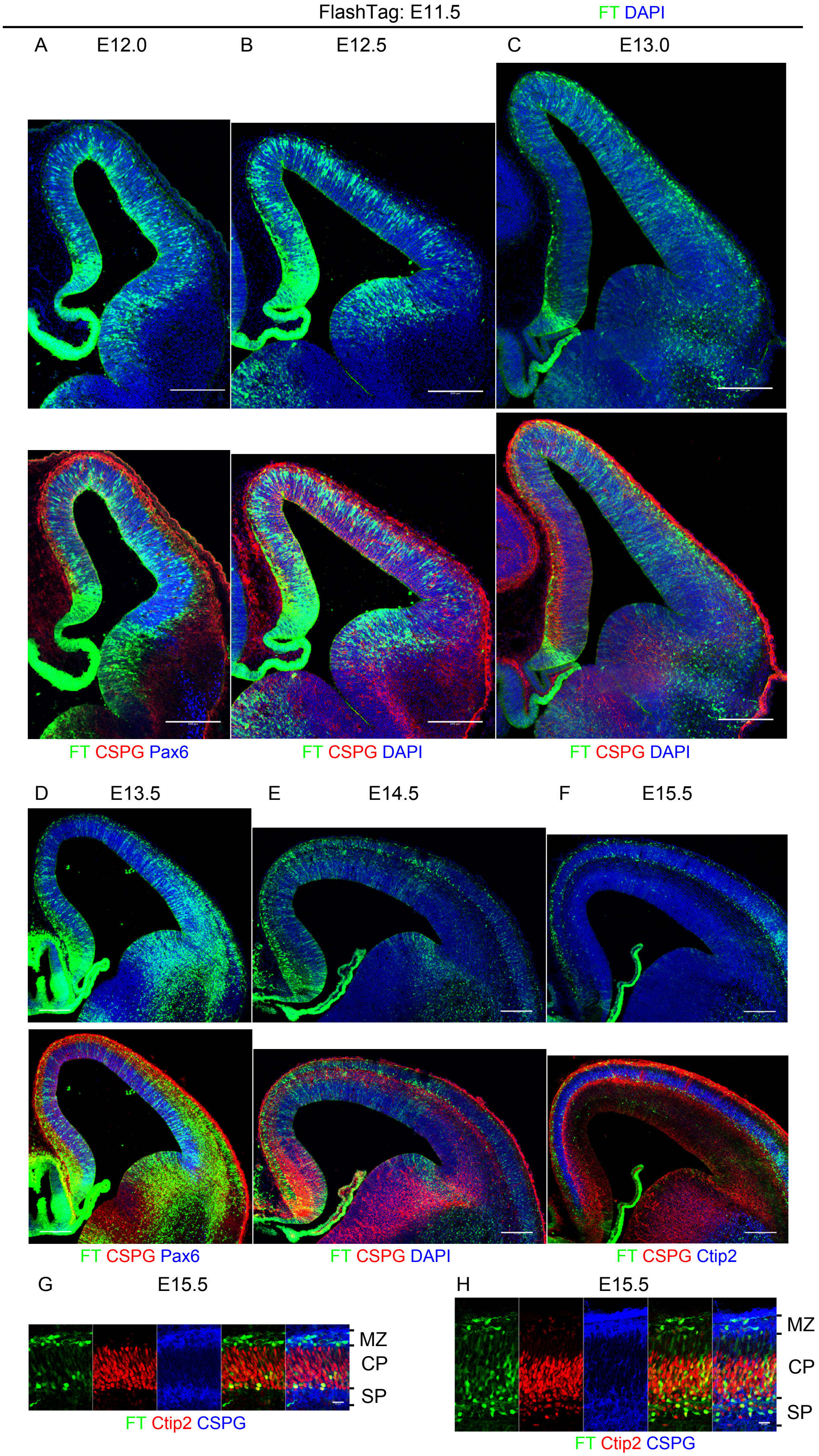

### Figure S4

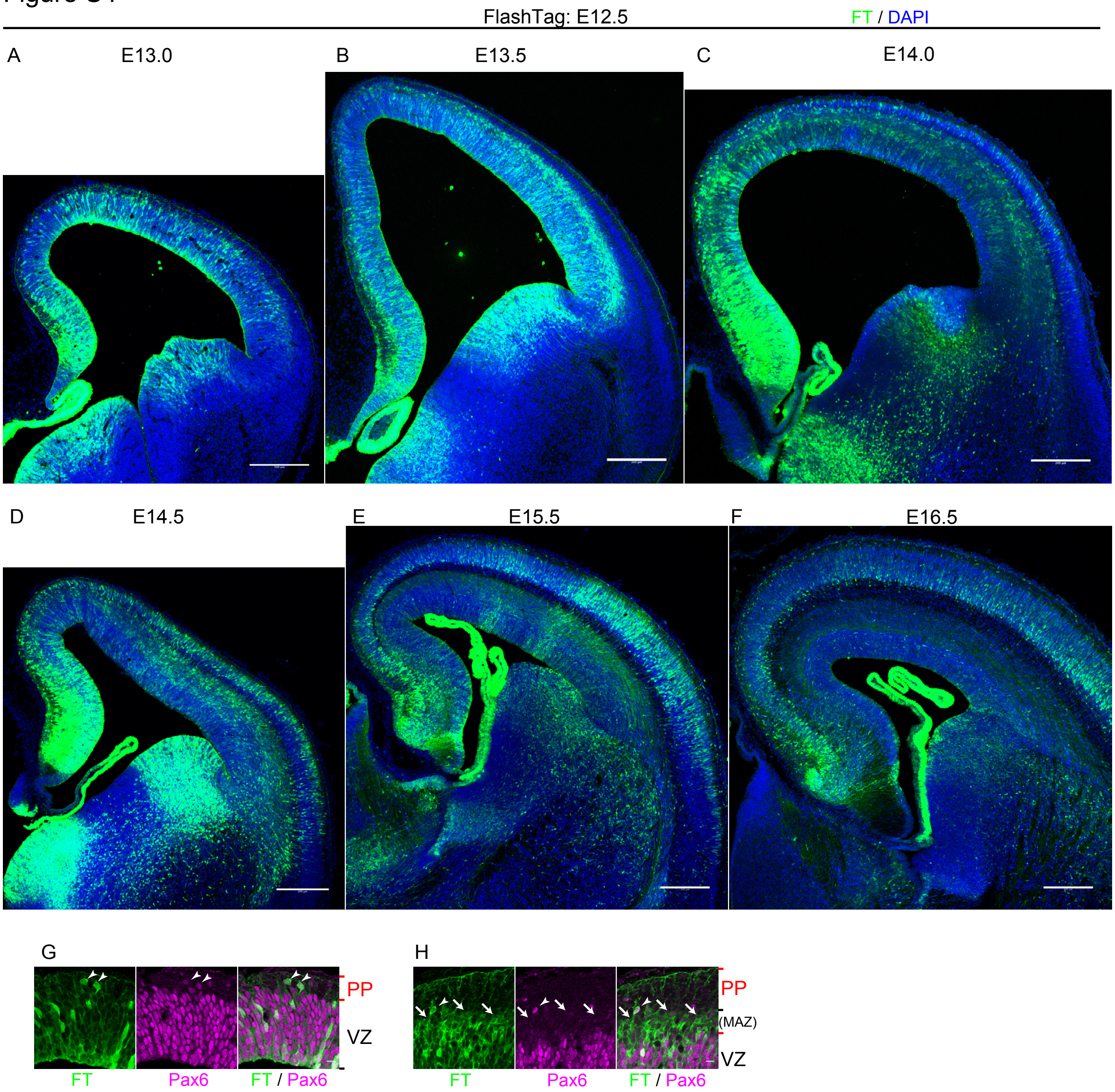

### Figure S5

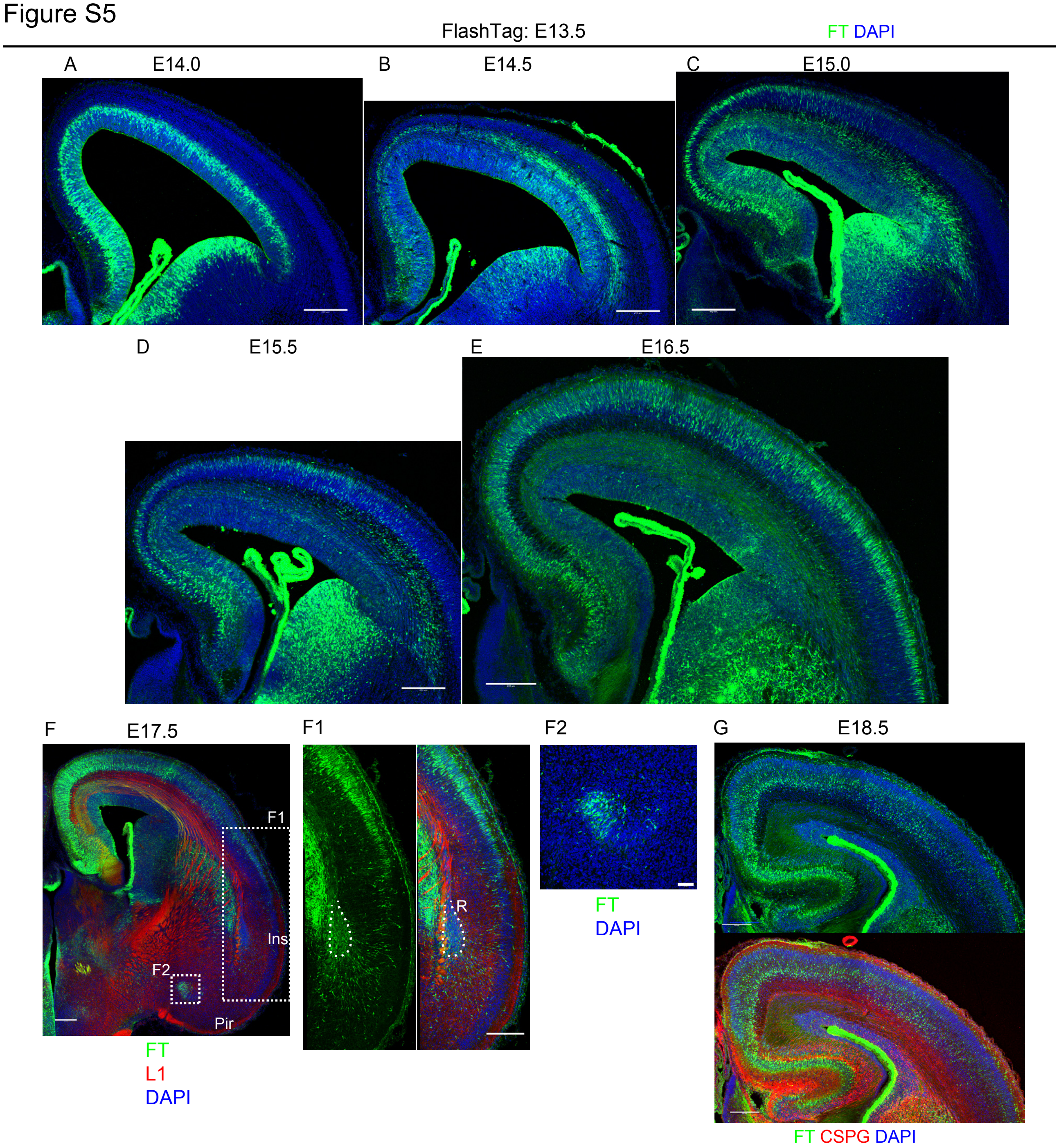

### Figure S6

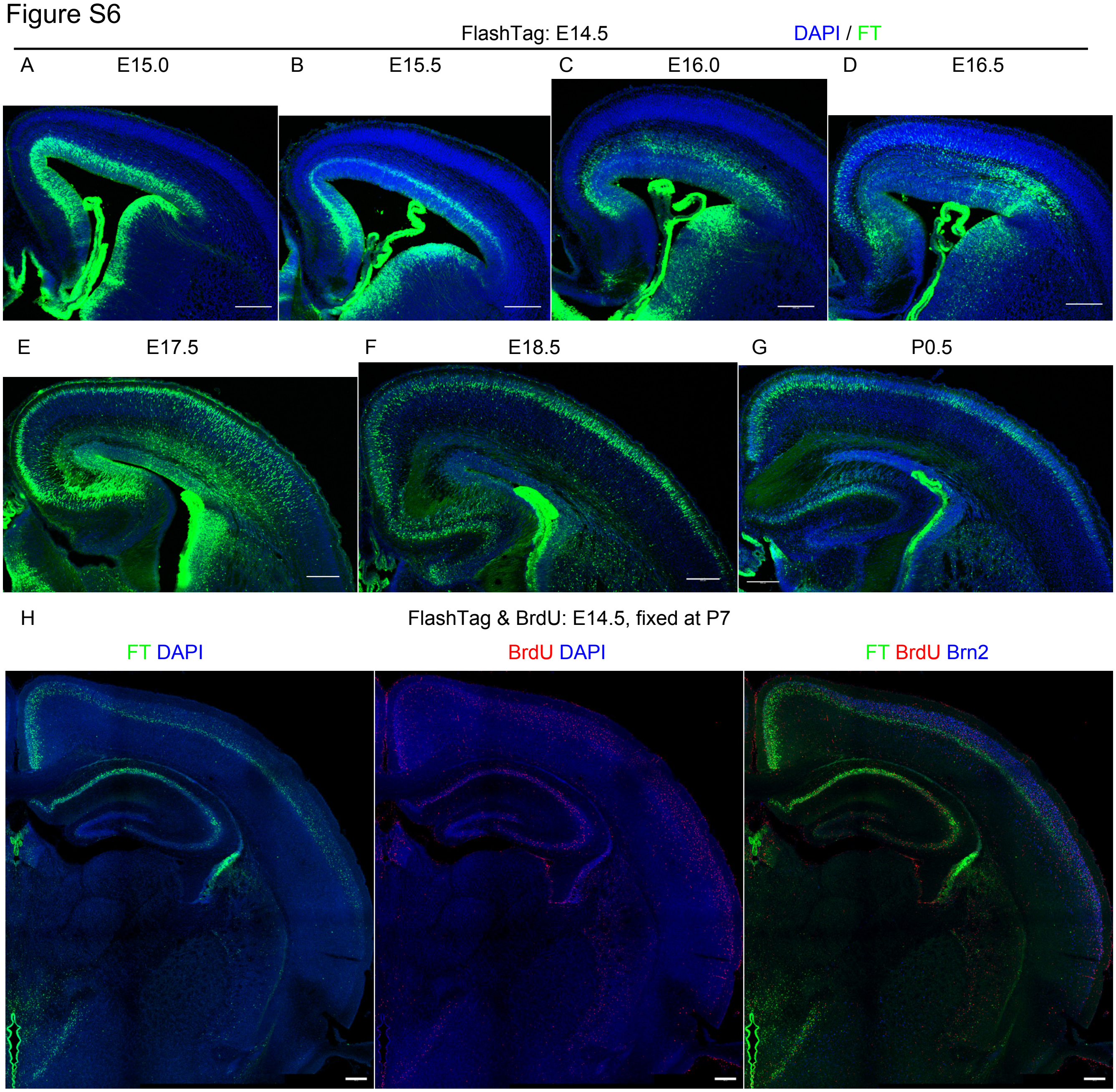

### Figure S7-1

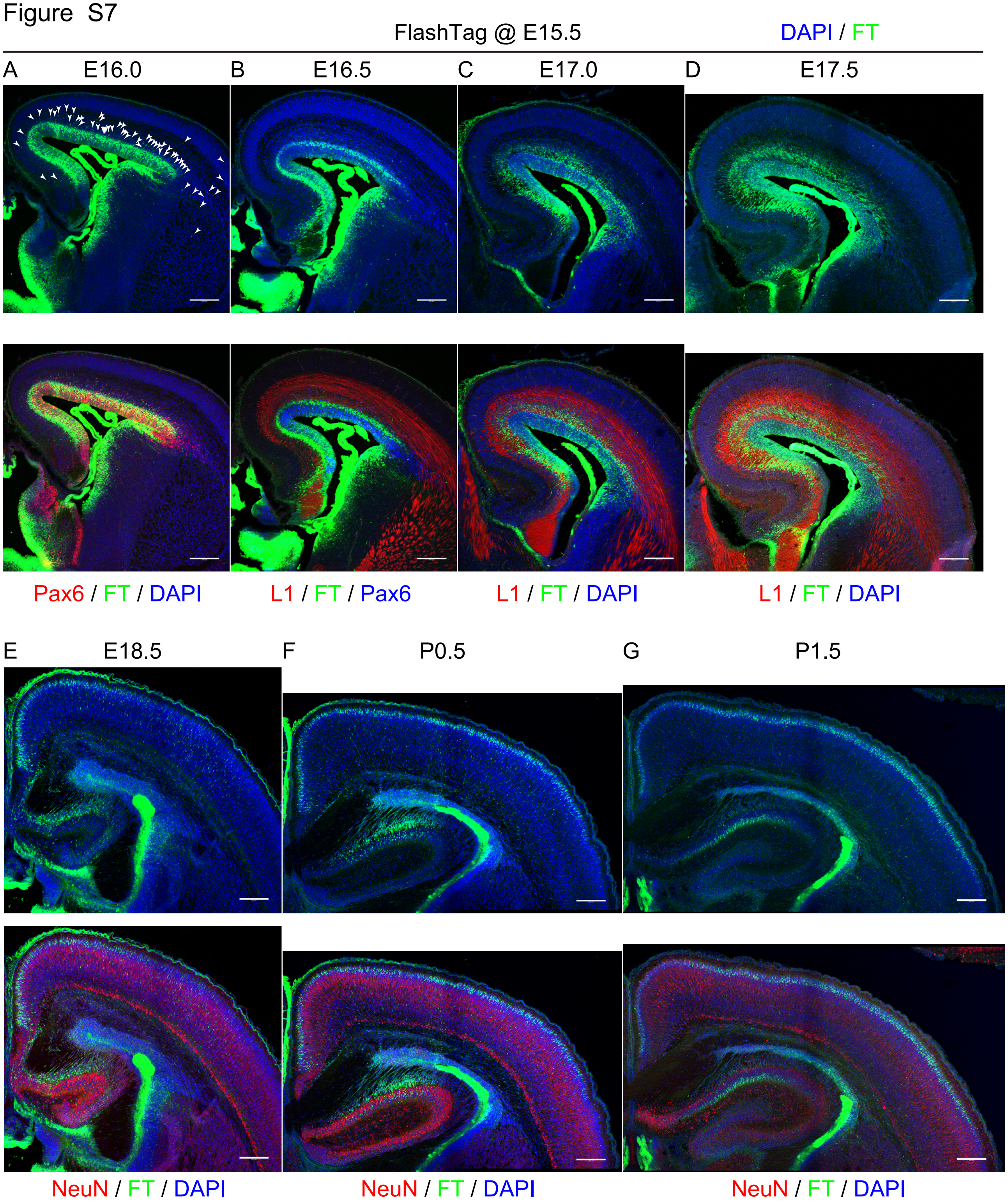

### Figure S7-2

Figure S7 (continued)

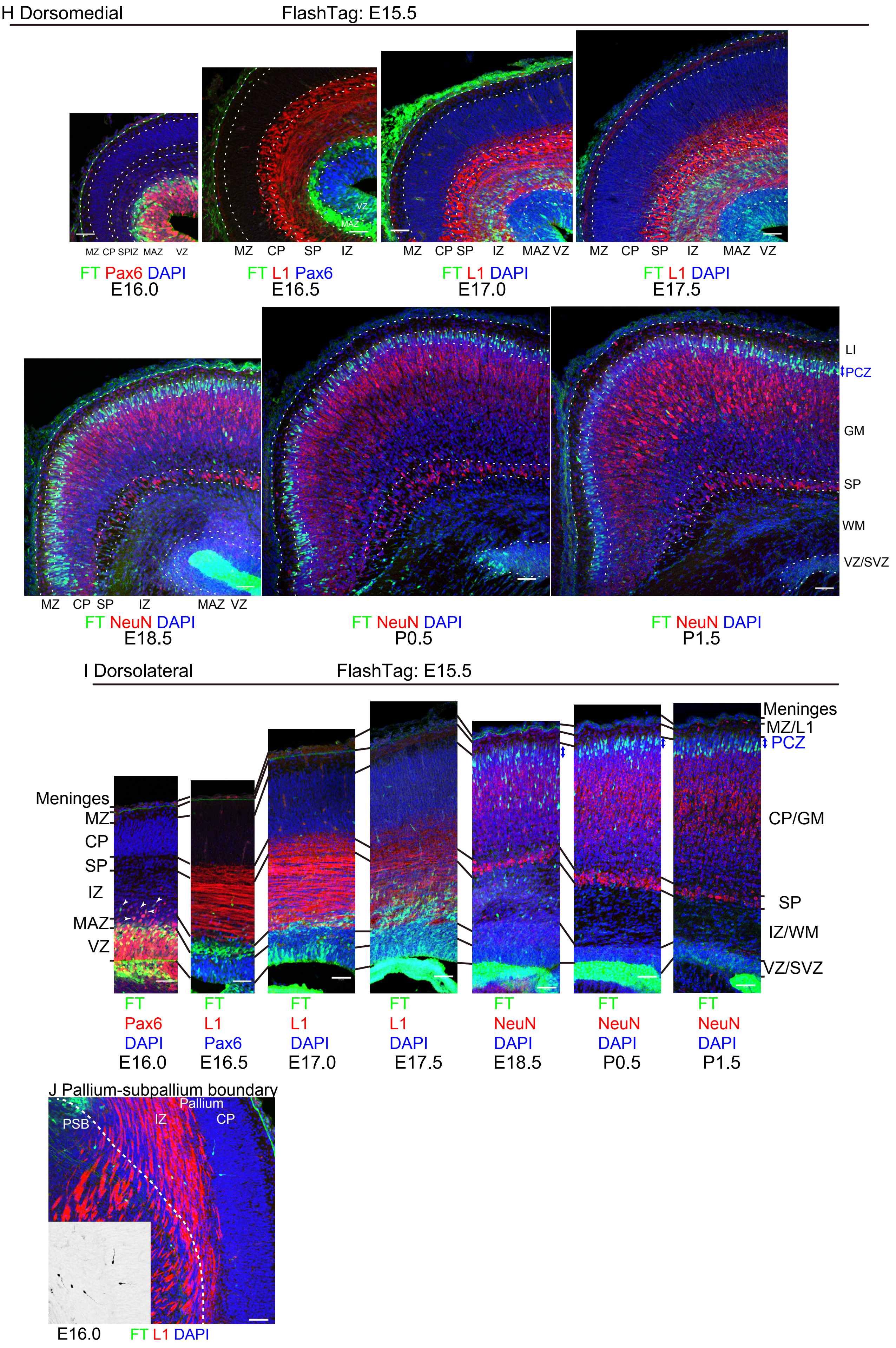

### Figure S8

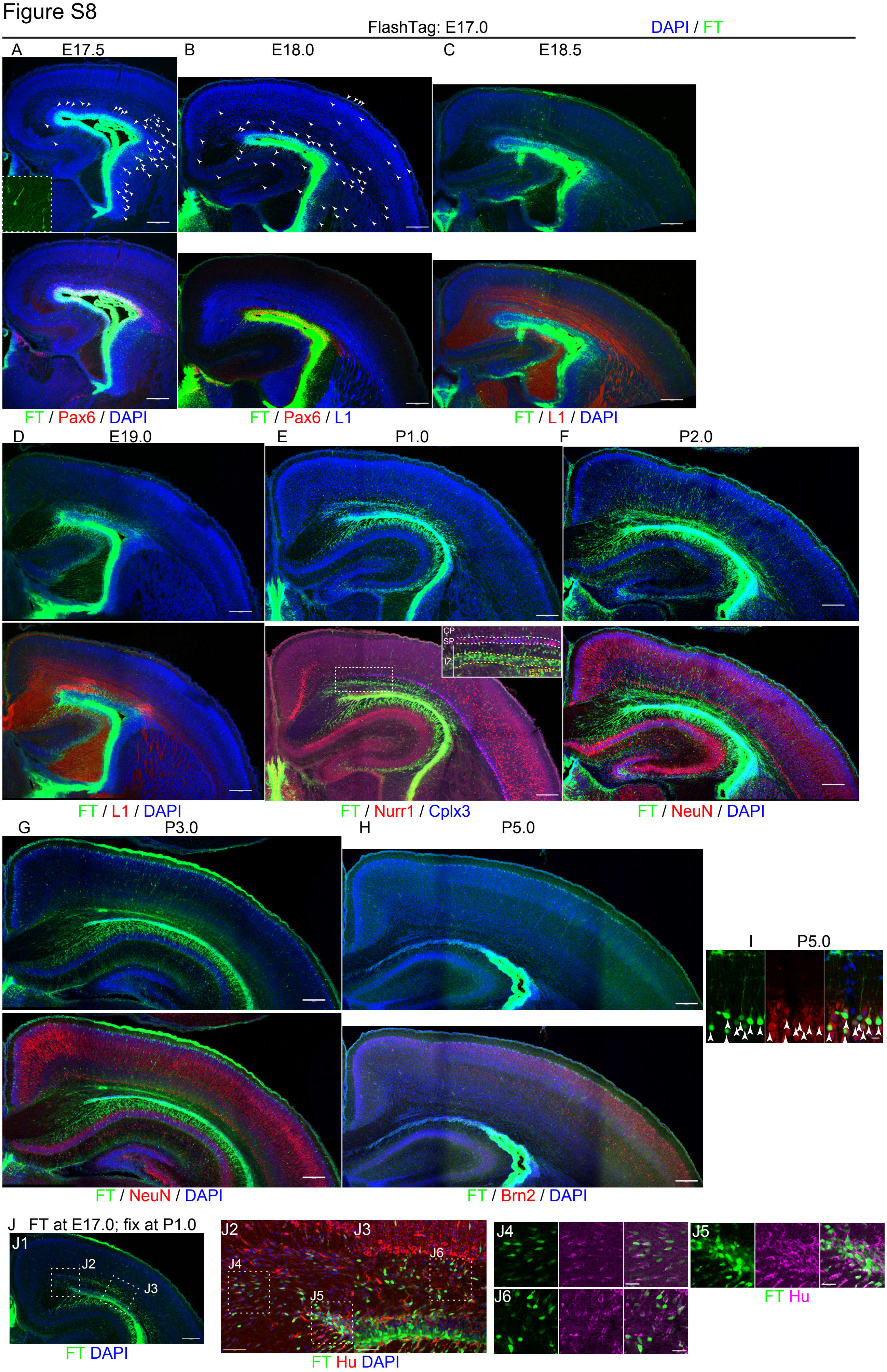

### Figure S9

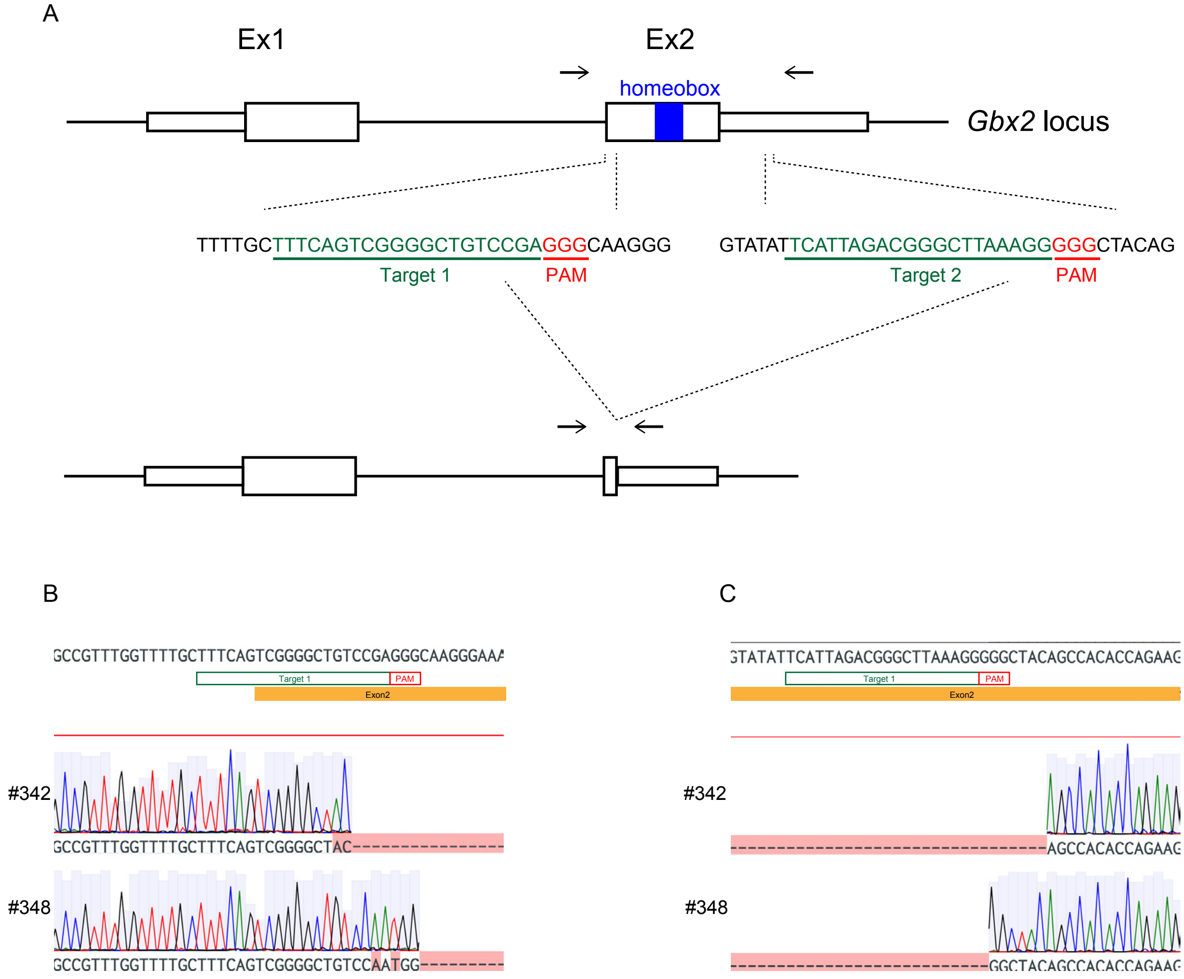
